# Integration spots organize genome plasticity and shape accessory-genome architecture in Ralstonia solanacearum

**DOI:** 10.64898/2026.02.24.707731

**Authors:** Upalabdha Dey, Jyotishpal Deka, Prerona Sharma, Mohit Yadav, Siddhartha Shankar Satapathy, Suvendra Kumar Ray, Aditya Kumar

## Abstract

The Ralstonia solanacearum species complex (RSSC) is a highly diverse plant pathogen whose evolution is shaped by horizontal gene transfer. We generated a complete, gap-free hybrid genome assembly of F1C1, a South Asian Phylotype I strain classified as R. pseudosolanacearum. The closed assembly resolves a bipartite genome (3.73 Mb chromosome; 2.03 Mb megaplasmid), enabling precise localization of mobile genetic elements. Using F1C1 together with 142 complete RSSC genomes, we implemented a lineage-stratified pangenome framework that reveals a hierarchically structured gene repertoire and shows that accessory gene content can discriminate host-associated lineages beyond core-genome. Pangenome-informed mapping of genome plasticity identified 651 conserved integration spots that concentrate accessory turnover and are enriched for adaptive functions, including type III secretion system effectors and antiviral defense systems (e.g., Wadjet and CRISPR-Cas). Genes within these spots display elevated *K*_*a*_/*K*_*s*_ relative to housekeeping functions, consistent with conflict-driven diversification and/or relaxed constraint. Together, these results link RSSC genome architecture to adaptive potential and provide a spot-based framework for genomic surveillance and resistance breeding in bacterial wilt pathosystems.

## 2 Introduction

The Ralstonia solanacearum species complex (RSSC) comprises a globally distributed group of soil-borne bacterial phytopathogens capable of infecting more than 300 plant species (Plant Health (EFSA PLH Panel) et al. 2019). Despite deep phylogenetic diversity spanning three species (*R. solanacearum, R. pseudosolanacearum, R. syzygii*) and four major phylotypes (Safni et al. 2014), RSSC members share a highly successful vascular wilt lifestyle (Genin and Denny 2012). RSSC strains persist in the rhizosphere, invade host roots, and spread through xylem vessels, ultimately producing vascular wilt and major yield losses in susceptible crops (An and Zhang 2024). Their broad host range, environmental persistence, and pronounced phenotypic variability have therefore made the RSSC a key model for understanding how bacterial pathogens diversify and maintain virulence in complex soil and plant-associated ecosystems (Vailleau and Genin 2023).

Genomics has reshaped how RSSC diversity is delineated and interpreted (Genin and Boucher 2004). Average Nucleotide Identity (ANI), support resolution of the complex into distinct species groups, and phylogenomics clearifies relationships among Asian (Phylotype I), American (II), African (III), and Southeast Asian (IV) lineages(Safni et al. 2014). Beyond taxonomy, comparative genomics repeatedly points to the accessory genome as a major driver of phenotypic differentiation (Sharma et al. 2022). In contrast to relatively conserved core, accessory genes often occur as linked modules concentrated within regions of genomic plasticity, where horizontal gene transfer and mobile elements enable rapid functional turnover (Remenant et al. 2010; Vandecraen et al. 2017; Bazin et al. 2020; Gonçalves et al. 2020). These variable regions are frequently enriched for host-interaction and competitive traits (Gonçalves, Queiroz, and Santana 2020), including secretion-associated repertoires such as secretion systems (Gonçalves et al. 2020), consistent with a model in which niche expansion and host adaptation are facilitated by the acquisition, loss, and reshuffling of discrete gene blocks (Fernandes et al. 2025).

A recurring limitation is that many publicly available RSSC genomes remain fragmented drafts. Fragmentated genome further obscures the structural context of mobile genetic elements, collapses repeats, and blurs the boundaries of variable regions, complicating population-scale inference about genome organization. RSSC genomes encode abundant prophages (Greenrod et al. 2022; Gonçalves et al. 2021), insertion sequences (Castillo et al. 2020), and anti-phage defense systems (Castillo et al. 2020), yet the field still lacks a unified view of how these components are arranged across lineages and how they co-localize with other adaptive modules within the flexible and modular genome (Prokchorchik et al. 2020).

Genomic analysis have supported the acquisition and dissemination of accessory genetic elements across RSSC strains through horizontal gene transfer (HGT). Traditional approaches for identifying such elements primarily rely on the detection of putative genomic islands (GIs). While effective in certain contexts, conventional GI-based methods capture only a subset of the RSSC mobile genome, since majority of the variable genome are structurally mosaic, may lose distinct compositional signatures over time through amelioration, or consist of small insertions and rearrangements that fall below the resolution of standard prediction algorithms (Gonçalves, Queiroz, and Santana 2020; Ding et al. 2024).

In contrast, the broader concept of integration sites, defined as contiguous blocks of variable genes flanked by conserved genes offers a more inclusive way to capture genome plasticity. Unlike strict genomic island definitions, this framework also encompasses prophages, integrative and conjugative elements, and older insertions regions that may have lost their canonical GI hallmarks over time (Bazin et al. 2020; Gonçalves et al. 2020). While previous works (Kang et al. 2025; Ding et al. 2024) have suggested that RSSC mobile regions can act as reservoirs for effectors and metabolic versatility, it currently remains unresolved whether adaptive modules are distributed idiosyncratically across genome or whether they preferentially accumulate at a finite set of recurrent integration loci that can be mapped across strains using conserved flanking backbones. Specifically, these gaps are relevant in the ecological context, as RSSC strains inhabit dense microbial communities in rhizospehere: where inter-bacterial competition (Aoun et al. 2024) and interaction with bacteriophages (Yang et al. 2023; J. Wang et al. 2024) might impose strong selective pressures. These ecological forces are likely to shape both defense repertoires and virulence-associated traits (Silva Xavier et al. 2019; Coupat et al. 2008), potentially influencing how accessory modules are gained, maintained, or lost across lineages.

To address these gaps, we generated a closed hybrid genome assembly for F1C1, a *R. pseudosolanacearum* (Phylotype-I) strain isolated from chili in Northeast region of India. We then integrated this assembly with 142 publicly available complete RSSC genomes in a lineage-aware pangenome framework to map RGPs into conserved integration spots. This population-scale approach moves beyond isolated GI predictions and evaluates whether virulence determinants (including Type III secretion effectors), interference systems (Type VI), and anti-phage defense modules co-localize within recurrent plasticity hotspots, thereby linking genome architecture to adaptive repertoire evolution across the RSSC.

## 3 Results

### 3.1 Hybrid genome assembly and annotation establish a high-quality RSSC reference

We generated a hybrid genome assembly for the local RSSC isolate F1C1 using Illumina short reads along with Oxford Nanopore long reads to establish a complete, high-contiguity reference suitable for comparative and pangenomic analyses. This approach yielded a complete, contiguous reconstruction comprising two circular replicons: a chromosome (3.73 Mb) and megaplasmid (2.03 Mb), totaling 5.76 Mb with an average GC content of 67% (*Supplementary Table 2*).

Because assembly quality directly governs the detection of structural features like coding regions, mobile regions, and genomic islands, we systematically benchmarked three state-of-the-art hybrid assemblers: MaSuRCA, Unicycler, and Trycycler. While all three integrate short and long reads, their strategies differ: Trycycler is consensus-based across multiple independent assemblies, whereas Unicycler and MaSuRCA operate as pipeline-based assemblers with less manual reconciliation. QUAST analysis revealed clear differences in contiguity and structural resolution among the resultant assemblies (**Supplementary Figure 1; Supplementary Table 3**). Both Trycycler and MaSuRCA resolved the genome into two contigs corresponding to the characteristic chromosome and megaplasmid. In contrast, Unicycler produced a fragmented reconstruction comprising ∼190 contigs (with >180 contigs exceeding 1 kb) and exhibited an inflated total assembly length relative to the two-contig assemblies, consistent with unresolved repeat expansions (**Supplementary Table 3**). High read mapping rates (>97%) and high sequencing depth (> 800×) across all assemblies confirmed strong representation of the underlying sequencing data (**Supplementary Table 4**), suggesting that the principal differences among assemblers reflect contiguity and structural resolution rather than insufficient coverage.

To evaluate completeness beyond contiguity metrics, we assessed the assemblies using both broad (*bacteria_odb10*) and lineage-specific (*burkholderiales_odb10*) BUSCO marker sets. Our analysis indicate that assembler that require human intervention and experience, i.e., Trycycler provide the strongest performance, achieving the highest recovery of single-copy orthologs while minimizing fragmented markers. While, MaSuRCA retained a higher fraction of fragmented and missing BUSCO markers despite producing a similarly contiguous reconstruction of genome (*Supplementary Figure 2*; *Supplementary Table 5*). Independent assessment with CheckM2 further supported the quality of the Trycycler assembly, estimating 100% completeness, 0.21% contamination, and a coding density of 87.6% (*Supplementary Table 2*). Based on this combined assesments of contiguity and completeness, we selected the Trycycler assembly for subsequent analyses and annotated it with BAKTA, resulting in 4,969 predicted coding sequences (CDS). The bipartite organization and functional landscape was visualised using a Circos representation as shown in **Figure 1A**, highlighting tracks for GC content, GC skew, and COG functional distributions across both replicons. F1C1 chromosome shows the classic GC-skew pattern expected in an RSSC strain, while the megaplasmid has a more uneven GC content and shifting gene density.

**Figure 1.**
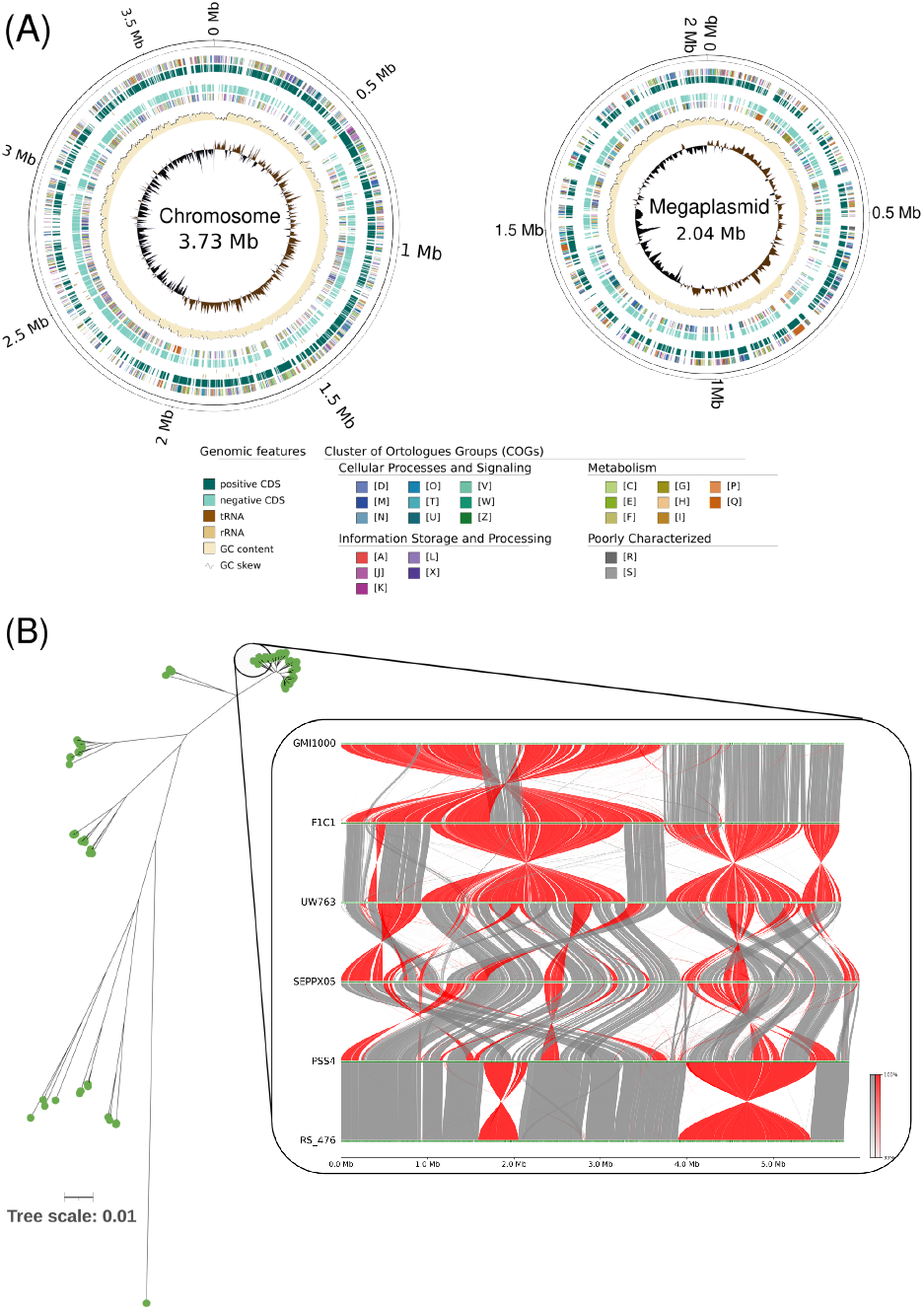
Complete genome assembly and phylogenetic placement of novel Ralstonia pseudosolanacearum strain F1C1. **(A)** Circos representation of the F1C1 genome illustrating the two-replicon architecture comprising the chromosome (3.73 Mb, *left*) and megaplasmid (2.04 Mb, *right*). Concentric tracks (inner to outer) show GC skew (black/brown indicating strand bias), GC content (cream indicating ∼67% average), and gene annotations on both strands with color-coded COG functional categories. The chromosome displays the expected bacterial GC-skew, whereas the megaplasmid exhibits more variable GC content and gene-density patterns typical of horizontally acquired genetic material. **(B)** Phylogenetic context and syntenic relationships of F1C1 among closely related RSSC strains. *Left*: distance tree derived from Mash k-mer distances showing the placement of F1C1 relative to phylogenetically related strains. *Right:* whole-genome alignment visualization using MUMmer comparing F1C1 with different strains (GMI1000, UW763, SEPPX05, PSS4, RS_476), where red segments indicate syntenic blocks and gray connectors highlight rearrangements and inversions. Overall, the alignment indicates strong synteny conservation across core regions while delineating variable segments likely associated with strain-specific adaptations and horizontally acquired elements.

Taxonomic placement of F1C1 strain was further assessed by using a Least Common Ancestor (LCA) approach based on genomic k-mer sketches (*k* = 31), which assigned F1C1 to the genus *Ralstonia* (family Burkholderiaceae), indicate its placement in the RSSC context (*Supplementary Table 1*). To further contextualize F1C1 genome variation against broader diversity, we performed an alignment free genome wide comparison of Mash distances. The neighbor-joining tree from placed F1C1 within the Phylotype-I cluster, grouping it with reference strains such as GMI1000 and UW763 (**Figure 1B**). Whole-genome synteny analysis against representative genome indicated extensive collinear blocks with high conservation and interrupted by localized rearrangements, inversions, and divergent segments. Together, these data support a model in which F1C1 retains a conserved RSSC backbone while concentrating strain-specific variation into genomic regions.

### 3.2 Core genome phylogeny reveals hierarchical population structure within the RSSC

To position F1C1 within the evolutionary context of the RSSC and to establish a robust framework for lineage-aware pangenomic inference, we curated a dataset of 142 complete genomes spanning a wide range of hosts and geographic origins. Pairwise Average Nucleotide Identity (ANI) comparisons separated these genomes into three species-level groups at the > 95% threshold, corresponding to established taxonomic assignments: *R. solanacearum, R. pseudosolanacearum*, and *R. syzygii*. Although ANI is well suited for species delimitation, it does not provide sufficient resolution to reconstruct within-species substructure required for interpreting gene-content evolution at population scale.

We therefore reconstructed a core-genome maximum-likelihood phylogeny using SNPs derived from an alignment of 2,756 single-copy orthologous gene families shared across all 142 genomes, and assessed node support using 1,000 ultrafast bootstraps (**Figure 2A**). To define nested population structure at multiple resolutions, we applied FastBAPS, which uses hierarchical Bayesian clustering. At the highest level, the primary clustering (L1) recapitulated the three ANI-defined species groups, supporting consistency between genome-wide similarity and phylogenetic signal. At finer resolutions, clusters aligned with the five phylotypes established by marker-based phylotyping (*egl, hrp, ITS*): Cluster 1 corresponds to phylotype IIB and IIC; Cluster 2 to I; Clusters 3 and 4 to III; and Cluster 5 to IV (**Figure 2A**). Integration of strain metadata further indicated that these phylogenetic and Bayesian partitions capture broad geographic and host-associated diversity while maintaining clear lineage coherence (**Figure 2B**).

**Figure 2.**
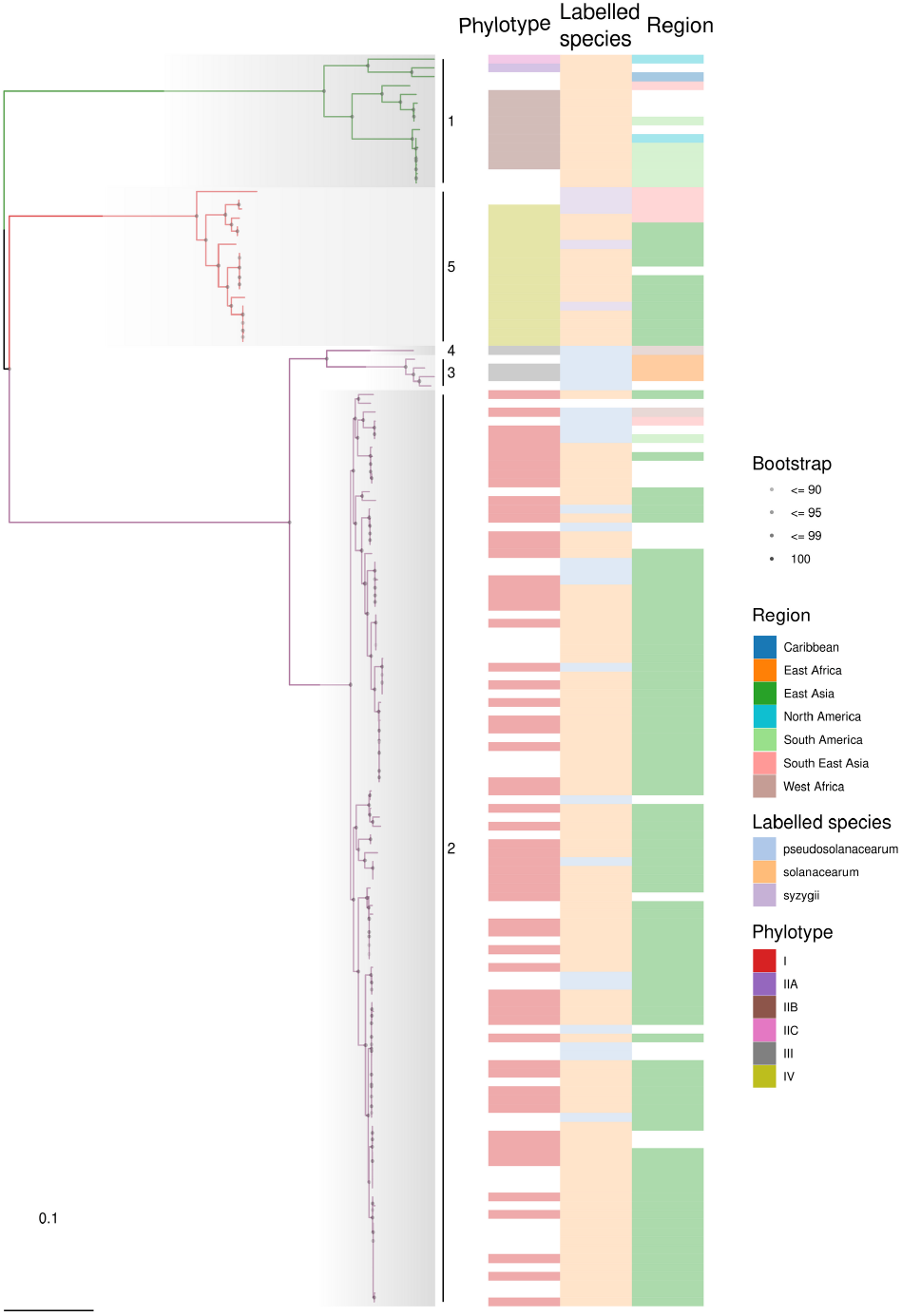
Core-genome phylogeny and geographic distribution of the RSSC. Maximum-likelihood phylogeny inferred from a concatenated core-gene alignment (3,263,944 bp from 3,348 genes) among 142 RSSC strains, summarizing the evolutionary relationships within the species complex. The phylogeny depicts the five major lineages that correspond to the defined phylotypes (I, IIA, IIB, III, and IV). The branch support is indicated by the use of bootstrap symbols (90: light gray, 95: medium gray, 99: dark gray, and 100: black circles). Metadata tracks provide geographic and taxonomic information. Phylotype labels show the hierarchical relationships at the population level, which transcend the species level. Species labels (*R. solanacearum, R. pseudosolanacearum*, and *R. syzygii*) indicate that the three recognized species are represented in the five phylotypes. The geographic labels show the overall bio-geographic distribution, with a preponderance of Phylotype IV in Asia and Phylotype II in the Americas.

Together, these analyses show that RSSC diversity is organized hierarchically, with deep species-level divisions and phylotype-level radiations. Current analysis provides an essential framework for downstream lineage stratified pangenomic analyses, allowing enumeration of genetic diversity and investigations of genomic plasticity to be interpreted through a population aware genomic context.

### 3.3 Lineage-stratified pangenome analysis reveals extensive functional plasticity and an open gene repertoire

Partitioning of RSSC pangenome into core versus accessory categories can be biased by uneven sampling of strains across clades. To mitigate this, we used a population-aware pangenomic framework that classifies gene families relative to inferred subclades (Horesh et al. 2021). This lineage-stratified strategy is especially important for structured species complexes like the RSSC because it separates genes that are genuinely rare across the entire population from genes that are stably maintained within particular lineages but appear rare under naïve global frequency thresholds.

Using high-resolution FastBAPS clustering (**Figure 2**; *Supplementary Figure 3A–B*), we first stratified gene families into core, intermediate, or rare categories within each lineage and then aggregated these into population-wide distribution classes (**Figure 3A**). Across the dataset, the RSSC pangenome contains 17,847 gene families, of which only 2,756 (∼ 15%) form the “Collection Core” conserved across all isolates (*Supplementary Figure 3B*). The remaining families fall into intermediate, rare, and lineage-restricted classes (**Figure 3B**), demonstrating that most RSSC genetic diversity lies within the accessory compartment.

**Figure 3.**
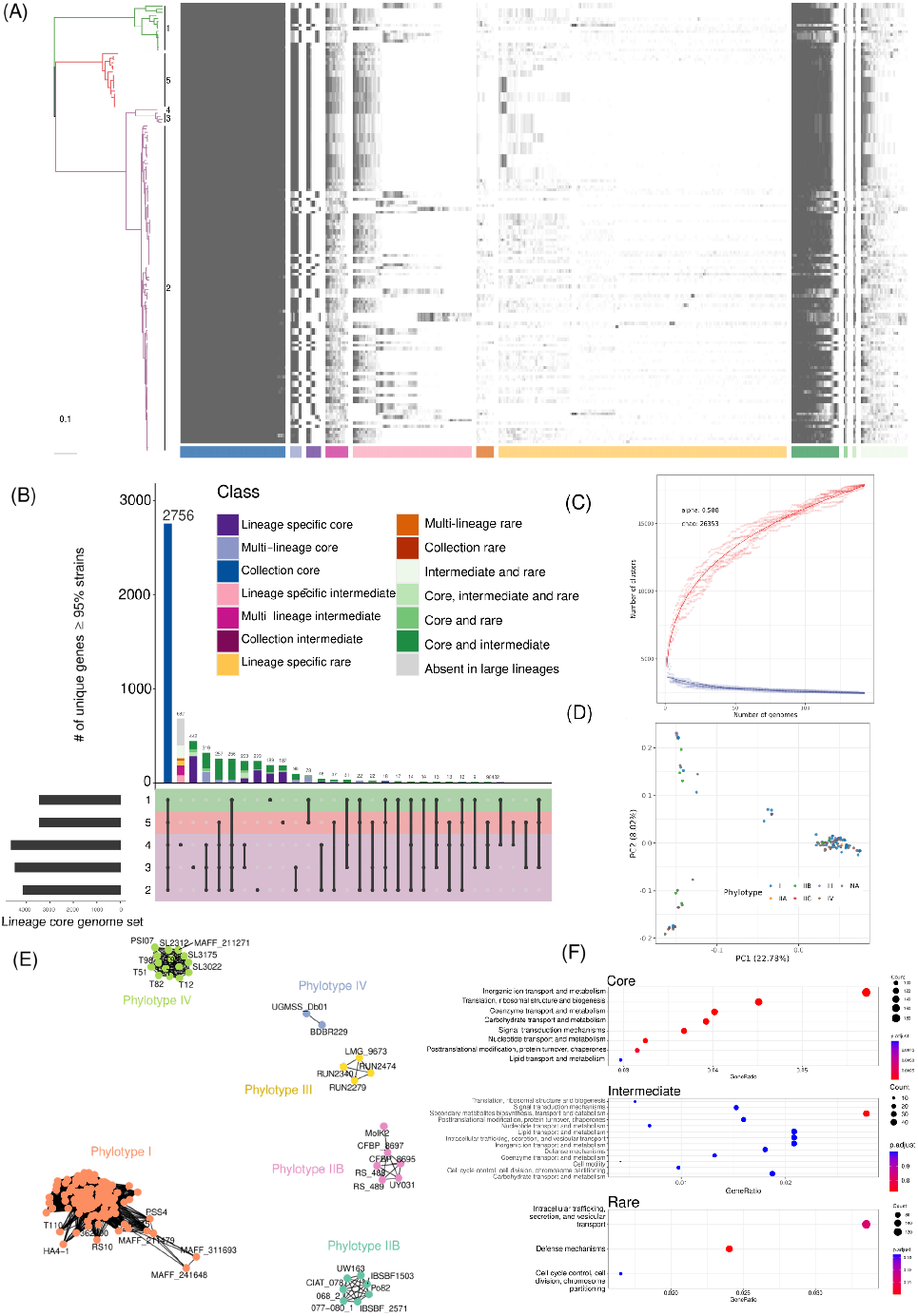
Pangenome structure and population organization of the RSSC. **(A)** Pangenome presence/absence matrix ordered according to Twilight classification and shown in conjunction with maximum likelihood phylogeny. The heatmap summarizes 17,847 gene families across 142 genomes, with columns organized according to pangenome partitions (core: dark; rare: light), highlighting the extent of the accessory genome. The accompanying dendrogram illustrates strong phylotype-level clustering with associated gene-presence patterns, as expected for lineage-specific accessory gene pools associated with ecological differentiation. **(B)** Gene accumulation rarefaction curve showing the open RSSC pangenome. The unsaturated curve and Heaps’ law alpha value (0.588, < 1) indicate ongoing gene discovery with increasing numbers of new genomes. Chao estimation of the pangenome size projects a total of approximately 26,353 gene families, reflecting high genetic diversity and ongoing horizontal gene transfer. **(C)** Principal component analysis of pangenome presence/absence data. PC1 and PC2 combined explain approximately 30% of the total variance. Isolates colored by phylotype show partial, but not complete, separation, indicating that while gene content patterns retain phylogenetic signal, extensive horizontal gene transfer precludes effective partitioning strictly on the basis of phylotype. **(D)** Network visualization based on Jaccard distances of the accessory genome, reflecting patterns of population structure complementary to the core genome phylogeny. Connected components (network clusters) organize isolates according to similar accessory genome content, illustrating that accessory genome similarity can extend beyond boundaries defined by core genome relationships, especially for phylotypes occupying similar ecological niches.

This lineage-aware framework also resolves the pangenomic “twilight zone”, namely genes whose frequencies differ across lineages and are therefore obscured by binary i.e., core versus accessory classifications. In particular, we identified a “Multi-lineage Core” comprising 335 gene families that behave as core genes within some clades while being absent from others (*Supplementary Figure 4*). Importantly, this heterogeneity varies strongly by lineage. Lineage 3 exhibits a substantially expanded core repertoire (median ∼330 families) compared to Lineages 1 and 2 (∼100–150 families) (*Supplementary Figure 3D–F*), suggesting that distinct evolutionary histories and ecological contexts have stabilized different lineage-associated gene pools.

Functional over-representation analysis showed that these partitions correspond to biological specialization. Lineage-specific core genes were significantly enriched for carbohydrate metabolism and secondary metabolite biosynthesis, consistent with niche associated adaptation. In contrast, lineage-rare genes were enriched for defense mechanisms, intracellular trafficking, and chromosome partitioning (**Figure 3F**; *Supplementary Figure 5*).

Rarefaction analyses further indicated a strictly open pangenome architecture: gene accumulation curves show no saturation across species or clusters (*Supplementary Figure 6*), with a Heaps’ law *α* = 0.58, consistent with ongoing gene discovery (**Figure 3C**). Principal component analysis (PCA) of gene content explained only ∼30% of the variance on the first two axes (**Figure 3D**), reflecting extensive horizontal gene transfer and turnover that limits simple low dimensional separation by gene content alone. Despite this plasticity, accessory gene composition retained a strong phylogenetic signal. Clustering based on Jaccard distances of accessory gene presence/absence recapitulated phylotype-level structure (*Supplementary Figure 7*). Notably, within Phylotype IIB, accessory content further resolved two distinct sub-clusters (**Figure 3E**), suggesting recent gene gain/loss events that may be associated with emerging host associated differentiation.

Collectively, these results define the RSSC pangenome as both open and structured. Much of the adaptive variation appears to be driven by modular acquisition and loss of accessory genes rather than by gradual divergence within the conserved core.

### 3.4 Compartment-specific *K*_*a*_/*K*_*s*_ patterns indicate reduced constraint in variable genome fractions enriched for mobility and defense functions

To characterize selective pressures across the RSSC pangenome, we estimated non-synonymous (*K*_*a*_) and synonymous (*K*_*s*_) substitution rates and summarized their ratio (*ω* = *K*_*a*_/*K*_*s*_) across the lineage-stratified gene classes. We restricted interpretation to gene families with defined rate estimates and treat elevated *ω* cautiously because it can be unstable for recently acquired genes with very low synonymous divergence (see Methods). At the global scale, purifying selection (*ω* < 1) predominates, affecting 70% of core gene families and 56% of accessory gene families. When stratified by lineage and gene-frequency class, *ω* shows a clear gradient that tracks population-wide conservation.

The Collection Core displayed the strongest constraint, with only 20% of gene families showing *ω* > 1 (**Figure 4A**). The proportion of *ω* > 1 increased in the multi-lineage core (23%) and peaked in the lineage-specific core (34%), indicating progressively reduced constraint as conservation becomes restricted to narrower phylogenetic subsets (**Figure 4B–C**). Functional enrichment supports a biological basis for this gradient. Highly constrained categories correspond to fundamental housekeeping processes (e.g., ribonucleoprotein complexes and aminoacid biosynthesis), whereas higher *ω* values concentrate in functions that interface with environmental variation, including transporters, kinases, and outer-membrane components, which may experience context-dependent divergence.

**Figure 4.**
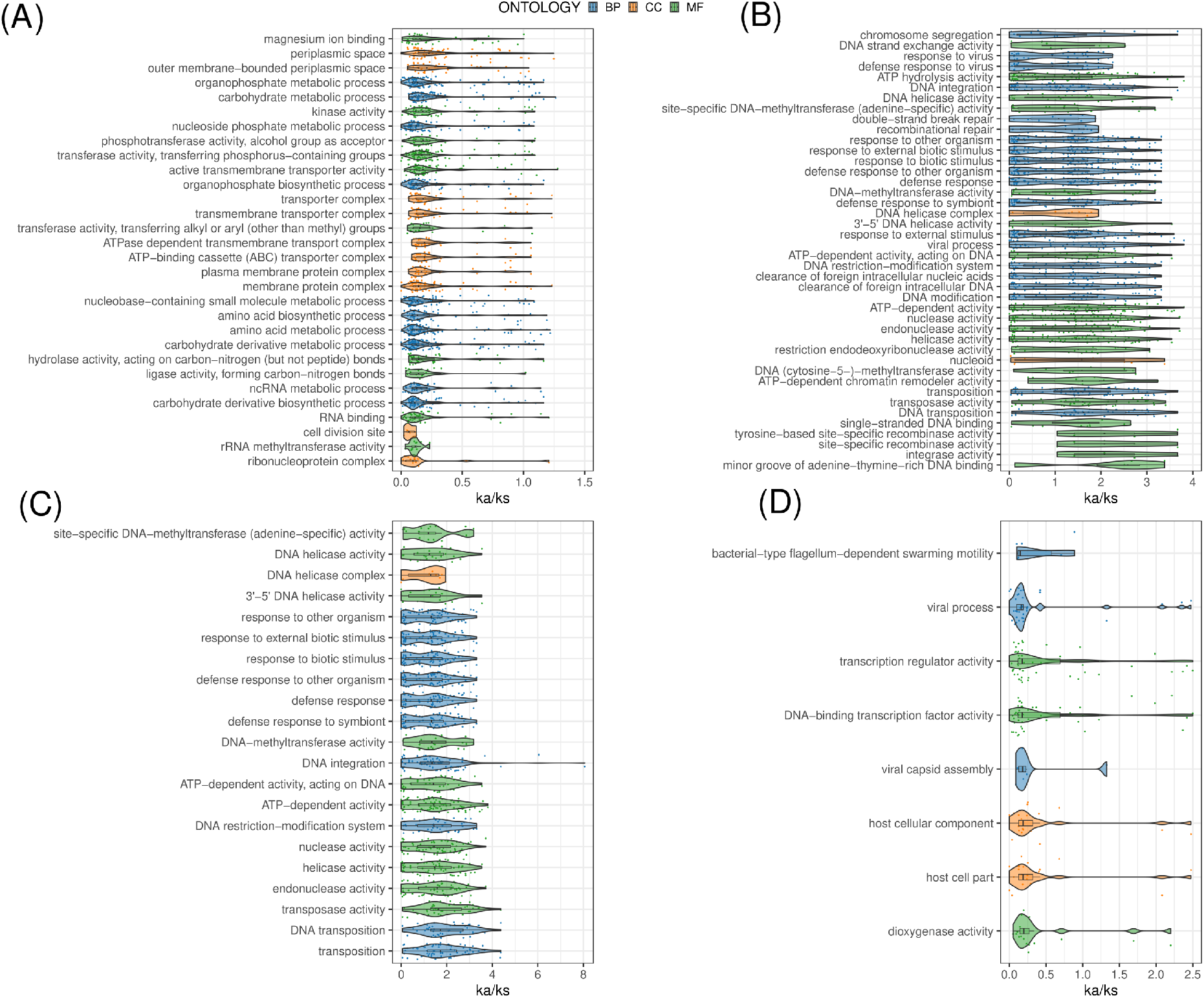
Evolutionary constraints on functional categories and pangenome compartments. Violin plots show Ka/Ks ratios for Gene Ontology categories in four pangenome compartments, emphasizing differences in evolutionary constraint for each compartment. Colors represent ontology terms: Biological Process (BP, blue), Cellular Component (CC, orange), and Molecular Function (MF, green). **(A)** Core genes are under strong purifying selection (Ka/Ks < 1) for essential functions like chromosome segregation, DNA repair, and metabolism, but have relatively higher Ka/Ks ratios for defense functions. **(B)** Rare genes have less stringent evolutionary constraints, with many functional categories having *K*_*a*_/*K*_*s*_ > 1, especially related to defense, DNA methylation, and transposition. **(C)** Lineage-specific core genes have intermediate levels of constraint, between the core and rare compartments. **(D)** Co-incident genes have variable levels of evolutionary constraint for functional categories, with defense and viral-related functions having higher evolutionary rates.

The pattern becomes most pronounced in highly variable accessory compartments. Gene families classified as “absent from major lineages” and “lineage-specific rare” showed high frequencies of *ω* > 1 (84% and 62%, respectively). These categories are enriched for mobility- and defense-related functions, including transposases and restriction–modification enzymes, consistent with reduced constraint and elevated divergence in mobile gene pools.

Because accessory genes can be gained or lost as linked modules, we tested whether the variable fractions in the RSSC pangenome with coordinated modularity. Identification of coincident accessory genes revealed a non-random modular structure (*Supplementary Figure 8*), with “Multi-lineage rare” and “Lineage-specific rare” families displaying statistically higher rate of co-association. Network reconstruction identified connected components spanning small clusters to larger modules. Notably, transcriptional regulators (including GntR, IclR, and DeoR families) within these modules frequently exhibited *ω* > 1, consistent with co-mobilization of regulatory elements alongside their targets, followed by divergence in new genomic contexts (**Figure 4D**; *Supplementary Figure 9–10*).

Overall, these results support a hierarchy of evolutionary constraint: core metabolic functions remain strongly conserved, whereas lineage-restricted accessory modules enriched in defense and mobility show reduced constraint and elevated divergence, consistent with modular gain/loss dynamics across the RSSC population.

### 3.5 Spatial organization of genome plasticity concentrates virulence, defense, and specialized metabolism in recurrent integration hotspots

To examine the spatial organization of RSSC genomic variability, we identified regions of genomic plasticity (RGPs) across all genomes and mapped them to conserved integration “spots” defined by flanking conserved genomic features (**Figure 5A**). This analysis shows that accessory evolution is embedded within a spatially constrained genomic context. Across the dataset, nearly 10,000 RGPs identified in the 142 complete genomes collapsed into 651 recurrent integration loci, indicating that accessory insertions are non-random and concentrated at discrete genomic sites (**Figure 5A**). We refer to loci containing >100 distinct gene families as “hotspots”, consistent with a higher capacity to acquire gene families repeatedly across isolates.

**Figure 5.**
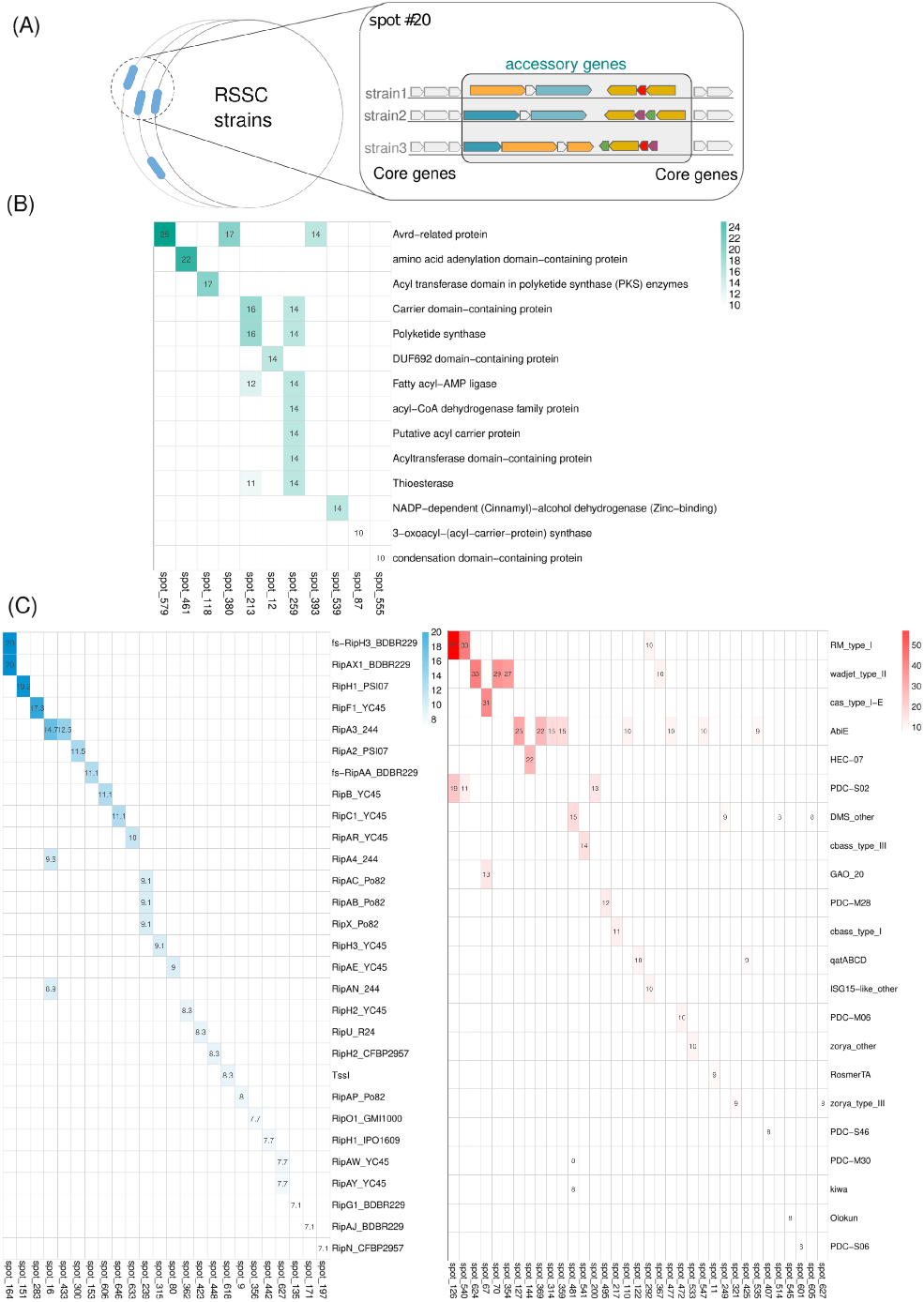
Functional structure of the accessory genome in regions of genomic plasticity. **(A)** Schematic representation of the dynamics of RGP integration in the RSSC pangenome. A total of 10,100 RGPs found in 142 genomes reduce to 651 distinct genomic spots, which correspond to hotspots of repeated integration of functionally related accessory genomic clusters. This functional structure is in line with the idea that genomic spots are specialized “docking stations” for horizontal gene transfer, which display stable chromosomal or plasmid localization propensities and are amenable to various gene content combinations. **(B)** Distribution of secondary metabolite biosynthetic gene clusters in major genomic spots. The bar diagram illustrates the non-random distribution of biosynthetic genes, which are repeatedly found in specific spots as preferred sites of integration. This functional structure implies that metabolic innovation is likely to be integrated into genomic contexts that are amenable to coordinated expression and regulation. **(C)** Functional specialization of genomic spots for virulence and defense. *Left:* Distribution of secretion system genes indicates that ∼50% of T3SS genes (5,495/10,853 total) are localized in specific spots, which display strong site preferences (e.g., *spot*#164 for RipH3, spot#16 for RipAN). *Right:* Defense system mapping reveals that 73% of PADLOC-identified defense genes (3,840 out of 4,870) are concentrated in distinct spots, which form “defense islands” rather than being randomly distributed. This functional structure implies that genomic spots have evolved to be specialized for hosting accessory functions, which can be assembled into complex phenotypes through horizontal gene transfer.

These integration spots display distinct architectural properties that reflect both replicon-specific constraints and differences in RGP length distributions. Chromosomal spots tend to accommodate larger gene blocks than megaplasmid spots (*Supplementary Figure 11*). Spot occupancy is also heterogeneous, ranging from broadly conserved loci shared across multiple lineages to restricted sites confined to specific clades (*Supplementary Figure 12*).

Functional mapping revealed that integration hotspots serve as recurrent platforms for distinct adaptive modules. Secondary metabolite biosynthetic gene clusters (BGCs) exhibited non-random clustering across major spots, with specific loci repeatedly functioning as preferred insertion sites for gene families relevant to metabolic pathways (**Figure 5B**). The preferential arrangement suggests that integration of BGCs might be constrained by local genomic contexts, which favor stable maintenance and coordinated regulation. For instance *spot*#579, *spot*#461 were enriched to Avrd-related protein, amino acid adenylation domain containing proteins (**Figure 5B**).

Virulence repertoires in RSSC displayed similarly structured pattern. Approximately 50% of RSSC type III secretion system (T3SS) effectors localized within RGP-associated spots, indicating that a large fraction of host-interaction machinery resides within hotspot-linked variable regions (**Figure 5C** and *Supplementary Figure 13*). Spot-effector associations were also specific: *spot*#164 showed enrichment for RipH3 families, while *spot*#16 repeatedly accumulated RipAN repertoires, indicating that effector repertoires are associated with recurrent genomic neighborhoods rather than being uniformly distributed. Principal component analysis of effector presence-absence patterns further showed that, despite modular assembly within spots, major repertoire differences remain phylogenetically structured, consistent with lineage-constrained horizontal transfer within a shared backbone scaffold(*Supplementary Figure 14*).

Interestingly, antiviral defense systems in RSSC showed a similarly strong association with genomic plasticity. Using PADLOC, we identified 4,870 defense genes spanning 116 distinct defense system types. We observed Wadjet-type-II, DMS, and zroya-type-III classes of defense systems are most prevalant contrast to others like RM-type-I, and cas-type-I (*Supplementary Figure 15*). A replicon stratified comparison further revealed specific localization preferences, with several defense families showing clear chromosomal versus non-chromosomal biases (*Supplementary Figure 16*). This pattern is consistent with a division between more stably maintained chromosomal defenses systems and more mobile defenses enriched on non-chromosomal replicons.

Most defense genes (73%; 3,840/4,870) localized within RGP-associated spots, forming defense-island like clusters across isolates. Defense integration was also highly structured at the spot level, i.e., spots with higher number of unique gene families usually have large number of unique defense gene families (*Supplementary Figure 17*). Moreover, several loci repeatedly accumulated high densities of defense genes across isolates, whereas other loci carried heterogeneous or lineage restricted defense complements. The distribution of defense repertoires across different spot and isolate summarizes the frequency of each defense system type within individual spots, revealing strongly non-random integration (*Supplementary Figure 18*). For example, Wadjet type II systems were repeatedly enriched at a small subset of integration spots (e.g., *spot*#624, *spot*#70, and *spot*#354), consistent with preferred “defense-island” sites (**Figure 5C (right panel)**). In contrast, other spots (e.g., *spot*#126) carried a range of defense repertoire types (including RM_type_I and PDC-S02), suggesting these loci function as more general defense integration hubs. The modular logic is exemplified by *spot*#81, where conserved flanking genes define a stable context while variable antiviral repertoires occupy the intervening segment as defense islands (*Supplementary Figure 19*).

Taken summarise, our findings point to a central organizing principle relevant to RSSC adaptive evolution: genes associated with adaptation are preferentially concentrated within recurrent integration hotspots. These integration sites likely to function as modular platforms where specialized metabolic pathways, virulence effector repertoires, and defense systems are assembled, reshuffled and might be helpful for HGT. Such spatial organization might facilitate diversification of adaptive traits while preserving the integrity of the conserved genomic organization through compartmentalized gene loss and gain.

## 4 Discussion

Our current study resents a complete, gap-free genome assembly of the South Asian Phylotype I RSSC isolate F1C1. We used this circular, closed reference to interpret population aware pangenome evolution at locus-level resolution. A long-read first hybrid assembly strategy, reconciled through the Trycycler consensus framework (Wick et al. 2021), was fundamental to resolve the bipartite genome architecture characteristic of the RSSC and to recover repetitive, insertion sequence prone regions that frequently fragment short-read assemblies in high-GC bacterial genomes. The final assembly comprises a 3.73 Mb chromosome and a 2.03 Mb megaplasmid, enabling unambiguous placement of repeats and mobilome that are otherwise difficult to resolve. Correct interpretation of genome plasticity needed correct genomic context, as the fragmented assemblies may obscure gene boundaries, misplace mobile genetic elements, or even generate the discrete genomic islands that reflect assembly artifacts without biological significance. By contrast, the closed F1C1 genome enables precise delineation and mapping of regions of genomic plasticity (RGPs) as discrete segments within defined replicons, thus providing a correct reference for comparative analysis in the evolutionary context.

The whole genome sequence of F1C1 also shows a functional compartmentalization that appears to be a hallmark of RSSC pathobiology (Genin and Denny 2012). This bipartite organization should not be considered as a strict functional separation but as a skewed representation of gene categories in replicons. However, the megaplasmid-mediated preponderance of adaptive and variable genes offers a functional platform through which the RSSC lineages can adapt phenotypically without compromising the integrity of the core cellular functions sustained in the chromosome. In F1C1, the essential cellular functions are encoded in the chromosome, while the megaplasmid is the main reservoir of strain-specific and adaptive genetic information. The two-component systems of RSSC also demonstrate the complexity of signaling transduction pathways that connect the detection of environmental cues to the control of virulence as a whole (Yadav et al. 2024), and how such complexity in regulation can facilitate adaptability in different environments.

At the population level, our lineage-stratified pangenome framework addresses a key limitation of the conventional core/accessory dichotomy in structured species complexes. Gene-family presence and frequency are not uniform across RSSC subclades, and global frequency thresholds can mis-classify genes that are stable within some lineages but absent from others. By defining gene conservation explicitly in terms of lineage-aware frequency classes, we resolve intermediate distributions that constitute the “twilight zone” of pangenomes. The identification of 335 gene families forming a “Multi-lineage Core,” conserved across multiple but not all lineages, demonstrates that RSSC pangenome evolution includes stable intermediate layers beyond a single universal core plus an undifferentiated accessory pool. We also observe heterogeneous retention of lineage-specific repertoires: Lineage 3 maintains a substantially larger stable accessory genome than Lineages 1 and 2, consistent with lineage-specific evolutionary pressures shaping gene-family retention and loss through differences in ecology, gene flux, and selective filtering. These patterns were identified through population structure analysis using hierarchical Bayesian clustering to partition our genomic dataset into evolutionarily coherent lineages.

Accessory gene content also appears to recapitulate recent ecological structure more strongly than core-genome phylogeny in specific comparisons. In our analyses, accessory variation provides superior resolution for host-associated clustering within Phylotype-IV, distinguishing Solanaceae- versus Musaceae-associated groups. This supports a model in which host association and host-expansion in RSSC populations can be mediated by horizontal acquisition and turnover of discrete “ecological gene modules”, rather than requiring gradual accumulation of adaptive mutations across the conserved core. This framing does not contradict core phylogeny; instead, it suggests that different genomic layers might encode different timescales and axes of diversification, with accessory genomic content often tracking recent eco-geographic sorting more sensitively than backbone variation.

Selective pressure analysis, based on Panaroo-defined orthologous gene families (Tonkin-Hill et al. 2020), further supports the existence of distinct evolutionary regimes in pangenome compartments. Analyses of regulators such as *PehR* and their structure and function support the evolution of transcriptional regulation to fine-tune pathogenicity responses (Yadav et al. 2023), providing a mechanistic explanation for the observed compartment-specific evolutionary patterns. In line, core housekeeping genes show strong purifying selection (*K*_*a*_/*K*_*s*_ < 1), indicative of evolutionary constraint on functions necessary for replication, gene expression, and central cellular physiology.

In contrast, genes involved in mobility, defense, and host interaction show enrichment for high Ka/Ks ratios, including values above unity (*K*_*a*_/*K*_*s*_ > 1), indicative of episodic selection and arms-race evolution at host-pathogen interfaces. This is most evident in lineage-specific rare compartments, where 84% of genes absent from large lineages show *K*_*a*_/*K*_*s*_ > 1. However, interpretation of *K*_*a*_/*K*_*s*_ ratios for low-frequency or recently acquired genes must be made with caution, as short divergence times, difficulties in alignment, and uncertainty in orthology may contribute to inflated *K*_*a*_/*K*_*s*_ ratios; hence, the most conservative interpretation is the compartment-level change in selection constraint.

One of the key findings of this study is that gene turnover is organized in space around conserved integration sites (Fernandes et al. 2025; Remenant et al. 2010). Using PanRGP (Bazin et al. 2020), we have identified 651 conserved integration sites, or “spots,” that concentrate genomic plasticity and are targeted by horizontally acquired modules.

These regions of the genome are highly enriched for systems that are central to bacterial defense and pathogenicity. Specifically, 73% of PADLOC-annotated defense genes (Payne et al. 2021), and 50% of Type III secretion system effectors, are found within spots. For instance, Wadjet systems are highly enriched at *spot*#624, while RipH3 effectors are preferentially associated with *spot*#164. This distribution, in aggregate, represents a pattern of locally constrained but statistically enriched genomic plasticity, wherein integration events are not randomly distributed across the genome but rather concentrated at specific loci that are apparently permissive to recombination and horizontal acquisition while avoiding the disruption of essential backbone functions. Such a modular organization of genome architecture provides a plausible mechanism for rapid adaptive evolution that is consistent with long-term genomic stability, wherein accessory genes that are co-present in these permissive regions can be swapped while the underlying architecture is comparatively conserved.

These results also have practical implications for the control of bacterial wilt in South Asia. High effector plasticity indicates that resistance strategies focused on highly variable effectors could be less long-term effective, as has been suggested by comparative studies indicating that highly virulent isolates often possess unique effector profiles (Kang et al. 2025). In this regard, factors identified through “multi-lineage core” analysis and other considerations that are conserved across lineages could represent more reliable targets for functional analysis and experimental validation.

In addition, the broad anti-phage defense profile evident in RSSC strains suggests that phage therapy strategies will likely need to be carefully tailored to match phage susceptibility profiles to bacterial defense profiles. Non-specific phage formulations could be less effective against strains possessing high-level modular defenses such as Wadjet and CRISPR-Cas systems. The discontinuous distribution of defense systems across lineages also suggests that clade-specific biocontrol strategies may be required.

In this regard, a genomic spot-informed surveillance strategy could provide practical utility. Analysis of the occupancy profiles of selected virulence and defense-associated integration spots in field populations may enable early warning of changes in epidemic potential or phage susceptibility, and thus provide support for more proactive and genomically informed disease management.

## 5 Methods

### 5.1 Culture and colony isolation of *Ralstonia pseudosolanacearum* F1C1

The highly virulent *R. pseudosolanacearum* strain F1C1 isolated from a wilted chili plant sampled from a field near Tezpur, Assam (Kumar et al. 2013). F1C1 glycerol stock was recovered from a deep-freeze storage and streaked onto a nutrient-rich solid medium (10 g/L peptone, 1 g/L yeast extract, 1 g/L casamino acids, 5 g/L glucose, 50 mg/L triphenyltetrazolium chloride (TTC), 15 g/L agar) under aseptic conditions, followed by incubation at 28°C for 48 hours.

F1C1 clones were recognized by their flat, irregular, fluidal colonies, which were pearly cream-white with a distinct pink center (Sarkar et al. 2025). The standard *R. pseudosolanacearum* F1C1 clones were streaked onto fresh nutrient-rich solid agar and maintained at 28°C for 48 hours. A single colony was transferred into a 15 ml culture tube with nutrient-rich medium without agar and TTC and cultured overnight at 28°C while shaking at 200 rpm. Microscopic verification of bacterial purity preceded all further experimental steps. The prepared F1C1 strain was subsequently utilized for genome sequencing.

### 5.2 DNA extraction and Quality control

Genomic DNA from *R. pseudosolanacearum* isolate F1C1 was extracted using a column-based DNA extraction kit, and the resulting DNA was subjected to a quality control (QC) workflow prior to downstream sequencing. Specifically, DNA integrity was checked by agarose gel electrophoresis, and DNA quantity/purity were evaluated using both NanoDrop spectrophotometry (NanoDrop: 59 ng/µl, A260/280 = 2.18, A260/230 = 0.04, measured volume 40 µl) and Qubit fluorometry (Qubit dsDNA HS Assay Kit, Thermo Fisher Scientific, USA; Qubit dsDNA: 7.5 ng/µl), ensuring the DNA met quality requirements for subsequent library preparation.

### 5.3 Library preparation and sequencing

In order to generate complete genome assembly, *R. pseudosolanacearum* genomic DNA was sequenced using Illumina short-read and Oxford Nanopore long-read technologies. 100 ng of genomic DNA was end-repaired using the NEBNext Ultra II End Repair Kit (New England Biolabs, MA, USA), followed by purification with 1X AMPure XP beads (Beckman Coulter, USA). DNA barcode adapter ligation was performed using NEB Blunt/TA Ligase Master Mix (New England Biolabs, MA, USA) and purified with 1X AMPure XP beads. Qubit-quantified, adapter-ligated DNA samples were barcoded using PCR amplification with LongAmp Taq 2X Master Mix (New England Biolabs, MA, USA) and purified with 1.6X AMPure XP beads (Beckman Coulter, USA). The prepared libraries were sequenced on the Oxford Nanopore MinION platform.

For short-read sequencing, mate pair library preparation was performed following the Nextera Mate Pair Gel Plus protocol (Illumina Nextera Mate Pair library preparation guide Cat# FC-132-9001DOC, Part#15035209 Rev D). Approximately 4 *μg* of Qubit-quantified high molecular weight genomic DNA was subjected to tagmentation using the Nextera Mate Pair Sample Preparation Kit (FC-132-1001), whereby the genomic DNA was fragmented and simultaneously attached to Illumina adapters required for sequencing. The tagmented sample was cleaned up using HighPrep™ PCR beads (Magbio, AC-60050) and subjected to strand displacement. The 3-5 kb size range of the strand displaced sample was selected using 0.6% agarose gel electrophoresis, followed by gel extraction using Zymoclean Large Fragment DNA recovery kit (Zymo research, D4045). Size-selected DNA was circularized overnight using circularization buffer and ligase, followed by linear DNA digestion with DNA exonuclease to remove uncircularized molecules. The circularized DNA molecules were sheared using Covaris S220 ultrasonication to obtain fragments in the size range of 300-1000 bp. Sheared DNA was subjected to bead binding with Dynabeads® M-280 Streptavidin (Invitrogen, 11205D) to isolate biotinylated molecules. End repair, A-tailing, and adapter ligation were performed on the bead-DNA complex using components from the Nextera Mate Pair kit. The adapter-ligated sample was amplified for 15 cycles of PCR followed by HighPrep bead cleanup. The prepared mate pair library was quantified using Qubit dsDNA HS Assay Kit and validated for quality using High Sensitivity TapeStation (Agilent, reagents 5067-5585, tapes 5067-5584). The final library exhibited an expected size distribution of 300-900 bp with effective insert size ranging from 180-780 bp flanked by adapters of ∼120 bp. The normalized library (2.43 nM by qPCR) was loaded onto an Illumina flow cell for paired-end sequencing (2 × 150 bp) on the Illumina HiSeq platform following manufacturer’s protocols.

Illumina mate pair sequencing generated 2,599,374 paired-end short reads, while 59,758 raw reads were obtained from Oxford Nanopore long-read sequencing. FastQC v0.11.9 (https://github.com/s-andrews/FastQC) was used to assess adapter contamination and raw read quality (Andrews et al. 2012). Nextera 3’ adapter sequences were removed from Illumina mate pair libraries, followed by quality filtering with a Phred score (Q-score) cutoff of 30 using Trim Galore v0.6.7 (Krueger 2015). For Oxford Nanopore raw reads, Porechop (https://github.com/rrwick/Porechop) v0.2.4 was used for adapter trimming and quality control.

### 5.4 Hybrid de novo genome assembly and quality evaluation

#### 5.4.1 MaSuRCA assembly

Processed reads from Illumina paired-end and Oxford Nanopore sequencing were subjected to automated hybrid *de-novo* genome assembly using MaSuRCA v4.0.9 (Maryland Super-Read Celera Assembler). MaSuRCA (Zimin et al. 2013) integrates the advantages of overlap-layout-consensus (OLC) and de Bruijn graph approaches for genome assembly with limited computational resources. The FLYE_ASSEMBLY flag was enabled (`FLYE_ASSEMBLY=1`) in the MaSuRCA configuration file, with jellyfish hash size set to twenty times the estimated genome size (`JF_SIZE=114600000`) and CA_PARAMETERS error rate set to 0.25.

#### 5.4.2 Trycycler assembly

In addition to the F1C1 MaSuRCA assembly, a long-read first hybrid assembly approach was employed to generate an alternative genome assembly and benchmark the sequencing datasets (Wick et al. 2021; Wick, Judd, and Holt 2023). Due to recent advancements in single-molecule sequencing accuracy, bacterial genomes assembled primarily using long reads followed by short-read polishing have been reported to achieve highly accurate assemblies with fewer long-range structural errors compared to conventional short-read-first assembly approaches.

Adapter-trimmed Oxford Nanopore reads were further filtered to retain the top 90% of reads with the highest quality scores. Read length and other relevant statistics of the Oxford Nanopore library were calculated using NanoStat v1.6.0 (De Coster et al. 2018). Filtering lower-quality reads yielded a higher N50 value of approximately 12.7 kb with increased mean and median read lengths compared to adapter-trimmed sequences, though coverage was reduced from 50× to 45× (calculated using an estimated genome size of 5.7 Mb).

Oxford Nanopore reads representing the top 90% in quality were further subsampled into 12 semi-independent read subsets. These 12 sub-sampled read sets were assembled using three different long-read *de-novo* assemblers: Flye v2.9.1 (Kolmogorov et al. 2019), Miniasm v0.3 (H. Li 2016), and Raven v1.8.1 (Vaser and Šikić 2021). Each assembler was used with four different read subsets, generating 12 non-identical assemblies. All contigs from the 12 assemblies were clustered based on computed Mash distances (Ondov et al. 2016) using complete-linkage clustering and visualized as a dendrogram using FigTree (https://github.com/rambaut/figtree) v1.4.4 (Rambaut 2018). Five contigs from assemblies created by Flye (cluster G) and Miniasm (cluster K) formed three discrete clusters (3, 4, and 5) that were not comparable in length and were therefore discarded.

Clusters 1 and 2 comprised 10 and 12 contigs, respectively, with sizes comparable to chromosome and plasmid components of other *Ralstonia pseudosolanacearum* assemblies. Sequence coverage was consistent (∼46× for cluster 2 members and ∼44× for cluster 1 assemblies). Clusters 1 and 2 were selected as cogent representations of the chromosome and plasmid, respectively. Contigs from each cluster were reconciled, circularized, and oriented to begin with the same gene (dnaA or repA) for multiple sequence alignment using MUSCLE v3.8.1551 (Edgar 2004). Oxford Nanopore reads were distributed to corresponding clusters, and consensus sequences were generated for each cluster. The resulting consensus sequences were polished with Oxford Nanopore reads using Medaka v1.7.2 (https://github.com/nanoporetech/medaka), followed by Illumina reads using one round of Polypolish v0.5.0 (Wick and Holt 2022; Bouras et al. 2024), and POLCA v4.0.9 (Zimin and Salzberg 2020). Short-read polishing was expected to correct short-scale errors such as base substitutions and indels.

### 5.5 Assembly quality evaluation and comparison with other assemblies

We used Sourmash v3.5.0 (Pierce et al. 2019) to classify signatures created from the assembled F1C1 genome using the GenBank k=31 LCA database with `sourmash lca`. Sourmash classified the genome as an unspecified strain of *Ralstonia pseudosolanacearum*. Assembly quality evaluation, genomic statistics, and read recruitment were calculated using QUAST v5.2.0 (Gurevich et al. 2013), including metrics such as number of contigs, length, GC content, Nx and NGx values, genome fraction percentage, percentage mapped reads, and misassemblies. Additionally, QUAST was used to align and compare the F1C1 assembly to *Ralstonia pseudosolanacearum* GMI1000 as a reference.

Genome completeness was assessed based on expected single-copy orthologous gene counts using BUSCO (Simão et al. 2015; Seppey, Manni, and Zdobnov 2019) v5.4.3 with `bacteria_odb10` and `burkholderiales_odb10` lineage datasets. Additionally, CheckM v1.2.2 (Parks et al. 2015) lineage-specific workflow (`--lineage_wf`) was used to verify completeness and contamination. This workflow approximates the taxonomic resolution of the query genome, identifies clade-specific phylogenetically informative marker genes, and quantifies marker gene presence to estimate completeness and contamination levels.

### 5.6 Gene model prediction and functional annotation

The Bakta v1.8.2 pipeline (Schwengers et al. 2021) was used to annotate protein-coding sequences and non-coding genes in the assembled genome. Prodigal v2.6.3 was used for proteincoding gene prediction (Hyatt et al. 2010), while Aragorn v1.2.41 (Laslett and Canback 2004) and Barrnap v0.9 (https://github.com/tseemann/barrnap) were used to predict tRNA and rRNA genes, respectively. Bakta predicted protein sequences were functionally annotated using the eggNOG-mapper v2.1.9 pipeline (Cantalapiedra et al. 2021). Since eggNOG-mapper uses pre-computed phylogenetically clustered orthologs across the tree of life, it provides higher sensitivity for detecting seed orthologs and broader coverage compared to simple homology-based approaches (e.g., BLAST searches against smaller databases like SwissProt).

Cluster of Orthologous Groups (Galperin et al. 2019) (COG) were identified for each annotated protein-coding gene using COGclassifier (https://github.com/moshi4/COGclassifier), performing PSI-BLAST (Altschul et al. 1997) searches against the COG database (Galperin et al. 2025). Gene Ontology (GO) annotations were transferred from BLAST hits against SwissProt (Gasteiger, Jung, and Bairoch 2001) and TrEMBL (O’Donovan et al. 2002) databases using an in-house GO annotation pipeline.

### 5.7 Average nucleotide identity (ANI) calculation and comparative genomics

ANI is a sensitive *in-silico* method for delineating species boundaries (Jain et al. 2018), with 95% ANI typically considered the threshold for species delineation (Richter and Rosselló-Móra 2009). All-vs-all ANI calculations between RSSC complete assemblies and the F1C1 genome were performed using FastANI v1.33 (Jain et al. 2018), and visualized using UPGMA clustering with `seaborn` as implemented in ANIClustermap (http://github.com/moshi4/ANIclustermap). Mashtree v1.2.0 (Katz et al. 2019) was used to compare 142 RSSC genomes and create a distance-based phylogenetic tree, which was visualized using iTOL v6.7.4 (Letunic and Bork 2024). Closely related genome assemblies were compared using MUMmer v4.0.0 (Marçais et al. 2018), and inversions and syntenic regions were visualized using pyGenomeViz v0.4.2 (https://github.com/moshi4/pyGenomeViz).

### 5.8 Pangenome construction and population-informed partitioning

We queried the NCBI GenBank database for publicly available genome sequences and retrieved 158 strains belonging to the *Ralstonia* genus marked with “complete” assembly status using the NCBI datasets command-line interface. We retained 142 genomes encompassing strains from the Ralstonia solanacearum species complex (RSSC), specifically *R. solanacearum, R. pseudosolanacearum*, and *R. syzygii*.

For pangenome construction, all 142 genomes were uniformly annotated using the Bakta pipeline (Schwengers et al. 2021), followed by pangenome analysis using Panaroo v1.5.0 (Tonkin-Hill et al. 2020). The initial clustering step used a protein sequence similarity threshold of 95%. The `--clean-mode` option was set to ‘sensitive’, and MAFFT v7.505 was selected for aligning core genes present in at least 95% of samples. Pangenome accumulation curves and principal component analysis were performed using the `Pagoo R` package v0.3.2 (Ferrés and Iraola 2021), and the alpha parameter was estimated using the `micropan R` package v1.2 (Snipen and Liland 2015).

In addition to Panaroo’s default core/accessory genome partitioning scheme, we implemented a population structure-informed partition of the RSSC pangenome using `twilight` analysis (Horesh et al. 2021). Core and rare frequency thresholds were set to 0.95 and 0.15, respectively. Population groups were defined by hierarchical Bayesian clusters created using FastBAPS v1.0.8 (Tonkin-Hill et al. 2019).

### 5.9 Phylogenetic reconstruction and population structure analysis

Pairwise ANI values were calculated for all 142 RSSC genome assemblies using FastANI. Assemblies were filtered using a 95% ANI cutoff and clustered using a custom Python script (Harling-Lee et al. 2022) that calculates connected components using the `networkx` package (Hagberg, Swart, and Schult 2007). Clustered networks were visualized using Cytoscape v3.10.2 (Demchak et al. 2014). UPGMA clustering was performed using FastANI results and custom scripts implemented with the `pheatmap R` package.

For accurate RSSC phylogeny reconstruction, concatenated core gene alignments of 3,348 genes (3,263,944 bp) from Panaroo were filtered to retain only polymorphic sites using SNP-sites v2.5.1 (Page et al. 2016). The resulting alignment of 591,669 bp was used to construct a maximum-likelihood phylogeny using IQ-TREE v2.3.6 (Minh et al. 2020) with 1,000 ultra-fast bootstrap replicates and the GTR+F+I+R2 substitution model as suggested by ModelFinder(Kalyaanamoorthy et al. 2017). For accessory genome clustering, pairwise distances were calculated using scripts from the Grapple suite (Harling-Lee et al. 2022), results processed in `R` using the `igraph` package (Csardi and Nepusz 2006), and visualized using the `ggraph R` package (Pedersen et al. 2017).

### 5.10 Genomic plasticity regions, integration spots, and specialized gene systems

Accessory gene clusters (regions of genomic plasticity, RGPs) were detected across the RSSC pangenome using the PPanGGOLiN (Gautreau et al. 2020; Mainguy et al. 2023) PanRGP module v1.2.105 (Bazin et al. 2020). Integration spots and their characteristic features were merged with RGP data in R for statistical analysis (R Core Team 2021).

Antimicrobial resistance genes were detected using Abricate v1.0.1 (https://github.com/tseemann/abricate) against multiple databases like CARD (Alcock et al. 2020), VFDB (Liu et al. 2022). RSSC-specific Type III secretion system (T3SS) genes were identified using nucleotide sequences from the RalstoT3E database (Peeters et al. 2013), formatted as a custom Abricate database. Type VI secretion system (T6SS) genes were identified using experimentally validated sequences from the GMI1000 genome retrieved from the SecReT6 database (J. Li et al. 2015). Secondary metabolite gene clusters were identified using antiSMASH v7.0.0 (Blin et al. 2023) with Bakta-annotated GenBank files, using the `--taxon bacteria` parameter and `--minimal` flag.

Antiphage defence systems were identified using PADLOC (Payne et al. 2021) (Prokaryotic Antiviral Defence LOCator). PADLOC was run with the latest PADLOC-DB release and default detection thresholds and system-definition models (http://github.com/padloc/padloc). For coordinate consistent mapping to genome features, PADLOC was executed using per-strain protein FASTA and GFF3 annotations, and PADLOC outputs were retained for downstream analyses.

Abricate and PADLOC results were summarized and integrated with RGP spots in R based on strain identifiers and sequence coordinates from Bakta annotation. Virulence factors and secretion systems identified by Abricate, and defence systems identified by PADLOC, were mapped to genomic spots based on coordinate overlap, and per-spot statistics were calculated in R.

### 5.11 Ka/Ks estimation

Ka, Ks, and the selective pressure ratio (*ω* = *Ka*/*Ks*) were estimated for Panaroo defined gene families using KaKs_Calculator2 (D.-P. Wang et al., n.d.). For each gene family, a gene ID list was generated by extracting member identifiers across strains, and the corresponding protein sequences were retrieved from a concatenated FASTA of Bakta annotated proteins per-strain using seqtk (http://github.com/lh3/seqtk).

Multiple sequence alignments were constructed independently for each gene family at the protein level using MAFFT L-INS-i (invoked via mafft-linsi), which is appropriate for high-accuracy alignment of moderate-sized protein sets (Katoh et al. 2002).

To compute Ka/Ks, protein alignments were converted to codon-based nucleotide alignments using PAL2NAL (Suyama, Torrents, and Bork 2006), which back-translates amino-acid alignments onto corresponding CDS while preserving reading frame. Following codon alignment generation, duplicate sequences were removed at the nucleotide level using seqkit (Shen et al. 2016) to reduce redundancy and avoid unstable rate estimates driven by identical sequences.

Because KaKs Calculator operates on pairwise codon alignments in (quasi-)AXT format, the de-duplicated codon multi-FASTA alignments were converted into pairwise AXT blocks using a custom parser (http://github.com/kullrich/kakscalculator2). Pairwise Ka and Ks were then computed using kaksscalculator2 with the bacterial genetic code (-c 11) and the Nei–Gojobori (NG) method (`-m` *NG*), consistent with the analysis workflow. Output tables (Ka, Ks, *ω*) were subsequently parsed and summarized in R for downstream analyses.

### 5.12 Functional annotation and enrichment analysis

Functional annotation of the pangenome was performed using the `eggNOG-mapper` package. COG category annotations were parsed and processed in R for over-representation analysis of core, intermediate, and rare genes using the `enrichR` function from the `ClusterProfiler R` package v4.6.2 (Wu et al. 2021).

## 6 Data availibility

The genome assembly and raw sequencing data generated for Ralstonia pseudosolanacearum strain F1C1 have been deposited in the European Nucleotide Archive (ENA) under study accession PRJEB107717. The associated sample record is ERS28794515 (BioSample: SAMEA121249153). Raw sequencing reads are available as an Illumina NextSeq 500 mate-pair run (ERR16655914; experiment ERX16046286) and an Oxford Nanopore MinION run (ERR16656002; experiment ERX16046374). The assembly submission is registered as ERA35602558.

