## Supplementary file for "Integration spots organize genome plasticity and shape accessory-genome architecture in Ralstonia solanacearum"

### Supplementary Tables and Figures for Integration spots organize genome plasticity and shape accessory-genome architecture in Ralstonia solanacearum

Upalabdha Dey[1,*] Jyotishpal Deka[1] Prerona Sharma[1]
Mohit Yadav[1] Prof. Siddhartha Shankar Satapathy[2]

Prof. Suvendra Kumar Ray[1] Dr. Aditya Kumar[1,*]

2026-04-08

[1] Tezpur University, Molecular Biology and Biotechnology, Tezpur, Assam, 784028
[2] Tezpur University, Computer Science & Engineering, Tezpur, Assam, 784028

### Supplementary Tables

#### Sourmash classification of F1C1 isolate

Table 1: Sourmash lca classification for F1C1 with GenBank k=31 mer signature

| | |
|---|---|
| ID | contig_1 |
| status | found |
| superkingdom | Bacteria |
| phylum | Proteobacteria |
| class | Betaproteobacteria |
| order | Burkholderiales |
| family | Burkholderiaceae |
| genus | Ralstonia |

#### CheckM2 evaluation for contamination

Table 2: CheckM2 prediction of contamination and completeness status

| Name | F1C1 |
|---|---|
| Completeness_Specific | 100 |
| Contamination | 0.21 |
| Translation_Table_Used | 11 |
| Coding_Density | 0.876 |
| Contig_N50 | 3730463 |
| Average_Gene_Length | 338.6294957 |
| Genome_Size | 5765906 |
| GC_Content | 0.67 |
| Total_Coding_Sequences | 4977 |
| Total_Contigs | 2 |
| Max_Contig_Length | 3730463 |
| Additional_Notes | None |

#### QUAST assembly comparison

Table 3: QUAST assembly comparison report

| Assembly | MaSuRCA | Trycycler | Unicycler |
|---|---:|---:|---:|
| # contigs (>= 0 bp) | 2.00 | 2.00 | 190.00 |
| # contigs (>= 1000 bp) | 2.00 | 2.00 | 186.00 |
| # contigs (>= 5000 bp) | 2.00 | 2.00 | 45.00 |
| # contigs (>= 10000 bp) | 2.00 | 2.00 | 27.00 |
| # contigs (>= 25000 bp) | 2.00 | 2.00 | 17.00 |
| # contigs (>= 50000 bp) | 2.00 | 2.00 | 16.00 |
| Total length (>= 0 bp) | 5761817.00 | 5765906.00 | 6516656.00 |
| Total length (>= 1000 bp) | 5761817.00 | 5765906.00 | 6513801.00 |
| Total length (>= 5000 bp) | 5761817.00 | 5765906.00 | 6103199.00 |
| Total length (>= 10000 bp) | 5761817.00 | 5765906.00 | 5967296.00 |
| Total length (>= 25000 bp) | 5761817.00 | 5765906.00 | 5819634.00 |
| Total length (>= 50000 bp) | 5761817.00 | 5765906.00 | 5791475.00 |
| # contigs | 2.00 | 2.00 | 190.00 |
| Largest contig | 3727787.00 | 3730463.00 | 1500947.00 |
| Total length | 5761817.00 | 5765906.00 | 6516656.00 |
| GC (%) | 67.01 | 67.02 | 66.97 |
| N50 | 3727787.00 | 3730463.00 | 463151.00 |
| N90 | 2034030.00 | 2035443.00 | 18178.00 |
| auN | 3129858.90 | 3132098.00 | 778523.20 |
| L50 | 1.00 | 1.00 | 3.00 |
| L90 | 2.00 | 2.00 | 20.00 |
| # total reads | 36484198.00 | 36481365.00 | 36535382.00 |
| # left | 18255821.00 | 18254055.00 | 18280399.00 |
| # right | 18168619.00 | 18167552.00 | 18195225.00 |
| Mapped (%) | 97.43 | 97.46 | 100.49 |
| Properly paired (%) | 95.42 | 95.48 | 94.38 |
| Avg. coverage depth | 945.00 | 945.00 | 840.00 |
| Coverage >= 1x (%) | 100.00 | 100.00 | 99.73 |
| # N's per 100 kbp | 0.00 | 0.00 | 0.00 |

Table 4: QUAST assembly reads recruitment report

| Assembly | MaSuRCA | Trycycler | Unicycler |
|---|---|---|---|
| # total reads | 36484198.00 | 36481365.00 | 36535382.00 |
| # left | 18255821.00 | 18254055.00 | 18280399.00 |
| # right | 18168619.00 | 18167552.00 | 18195225.00 |
| # mapped | 35548169.00 | 35555951.00 | 36715845.00 |
| Mapped (%) | 97.43 | 97.46 | 100.49 |
| # properly paired | 34814232.00 | 34832812.00 | 34482202.00 |
| Properly paired (%) | 95.42 | 95.48 | 94.38 |
| # singletons | 13267.00 | 12691.00 | 185860.00 |
| Singletons (%) | 0.04 | 0.03 | 0.51 |
| # misjoint mates | 52678.00 | 45368.00 | 309134.00 |
| Misjoint mates (%) | 0.14 | 0.12 | 0.85 |
| Avg. coverage depth | 945.00 | 945.00 | 840.00 |
| Coverage >= 1x (%) | 100.00 | 100.00 | 99.73 |
| Coverage >= 5x (%) | 100.00 | 100.00 | 98.80 |
| Coverage >= 10x (%) | 100.00 | 100.00 | 98.55 |

#### BUSCO assembly completeness asessment

Table 5: BUSCO assembly completeness evaluation

| Assembler | BUSCO lineage | summary |
|---|---|---|
| Mausrca | bacteria_odb10 | C:97.6%[S:96.8%,D:0.8%],F:1.6%,M:0.8%,n:124 |
| Mausrca | burkholderiales_odb10 | C:96.3%[S:96.2%,D:0.1%],F:2.2%,M:1.5%,n:688 |
| Trycycler | bacteria_odb10 | C:100.0%[S:99.2%,D:0.8%],F:0.0%,M:0.0%,n:124 |
| Trycycler | burkholderiales_odb10 | C:99.7%[S:99.7%,D:0.0%],F:0.1%,M:0.2%,n:688 |
| Unicycler Bold | bacteria_odb10 | C:91.9%[S:88.7%,D:3.2%],F:6.5%,M:1.6%,n:124 |
| Unicycler Bold | burkholderiales_odb10 | C:91.3%[S:85.9%,D:5.4%],F:6.5%,M:2.2%,n:688 |

### Supplementary Figures

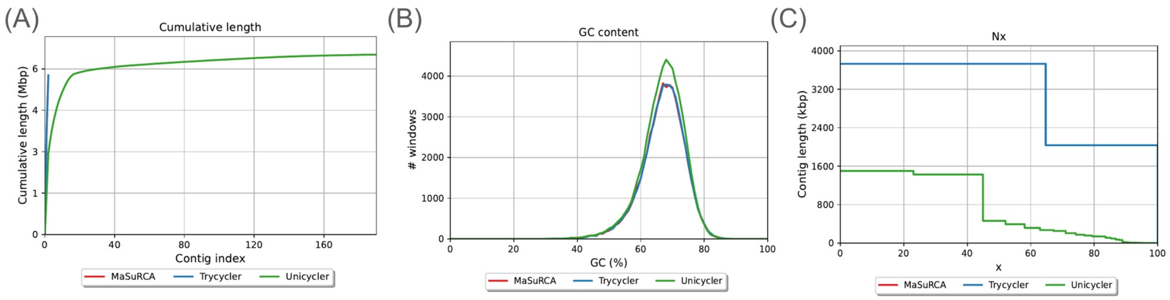

**Supplementary Figure 1. Comprehensive assembly quality assessment using QUAST.** Comparative evaluation of three genome assembly approaches (MaSuRCA, Trycycler, and Unicycler) for the F1C1 isolate. (A) Cumulative length plot demonstrates consistent total genome size (~5.7 Mb) across all assembly methods, confirming assembly completeness and validating the two-replicon structure (chromosome + megaplasmid). (B) GC content distribution analysis reveals consistent bimodal peaks across all assemblers, with the main peak at ~67% GC content characteristic of RSSC genomes, confirming assembly accuracy and lack of contamination. (C) Nx plot illustrating assembly contiguity metrics; the flat horizontal line for Trycycler indicates superior contiguity with only 2 contigs (N50 = N90), contrasting with the stepped decrease observed for Unicycler and MaSuRCA, which produced more fragmented assemblies with higher contig numbers.

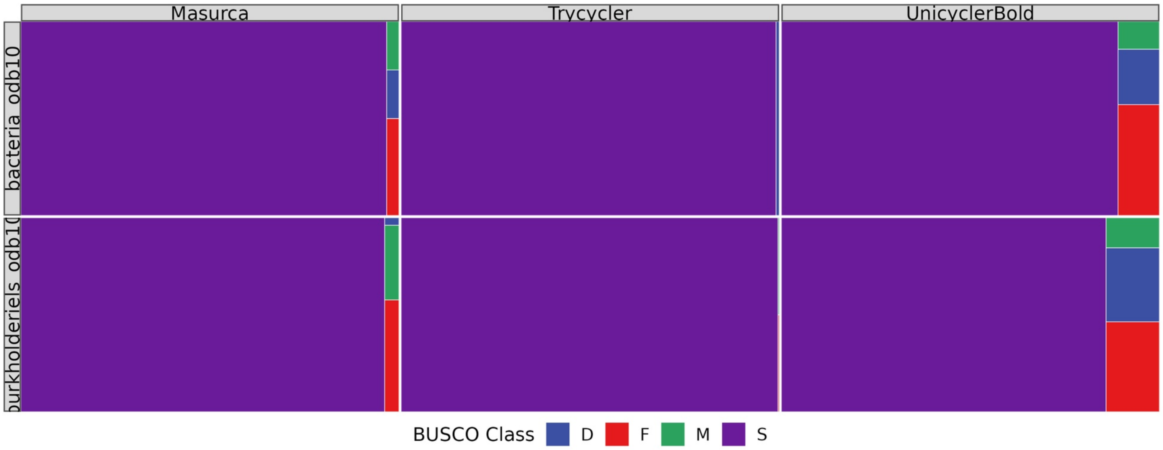
Masurca
Trycycler
UnicyclerBold
bacteria odb10
BUSCO Class
D
F
M
S

**Supplementary Figure 2. BUSCO completeness assessment across assembly methods.** Treemap visualization quantifying the recovery of phylogenetically conserved single-copy orthologs using both bacteria_odb10 and burkholderiales_odb10 lineage-specific marker sets. Each rectangle's area is proportional to the percentage of markers in each category: Complete Single-copy (dark blue), Complete Duplicated (light blue), Fragmented (orange), and Missing (red) genes. The Trycycler assembly (center) demonstrates superior performance with the highest proportion of complete single-copy markers and minimal fragmented/missing genes, particularly for Burkholderiales-specific markers, supporting its selection for downstream phylogenomic analyses. MaSuRCA and Unicycler show increased fragmentation rates, indicating assembly fragmentation affects gene model prediction quality.

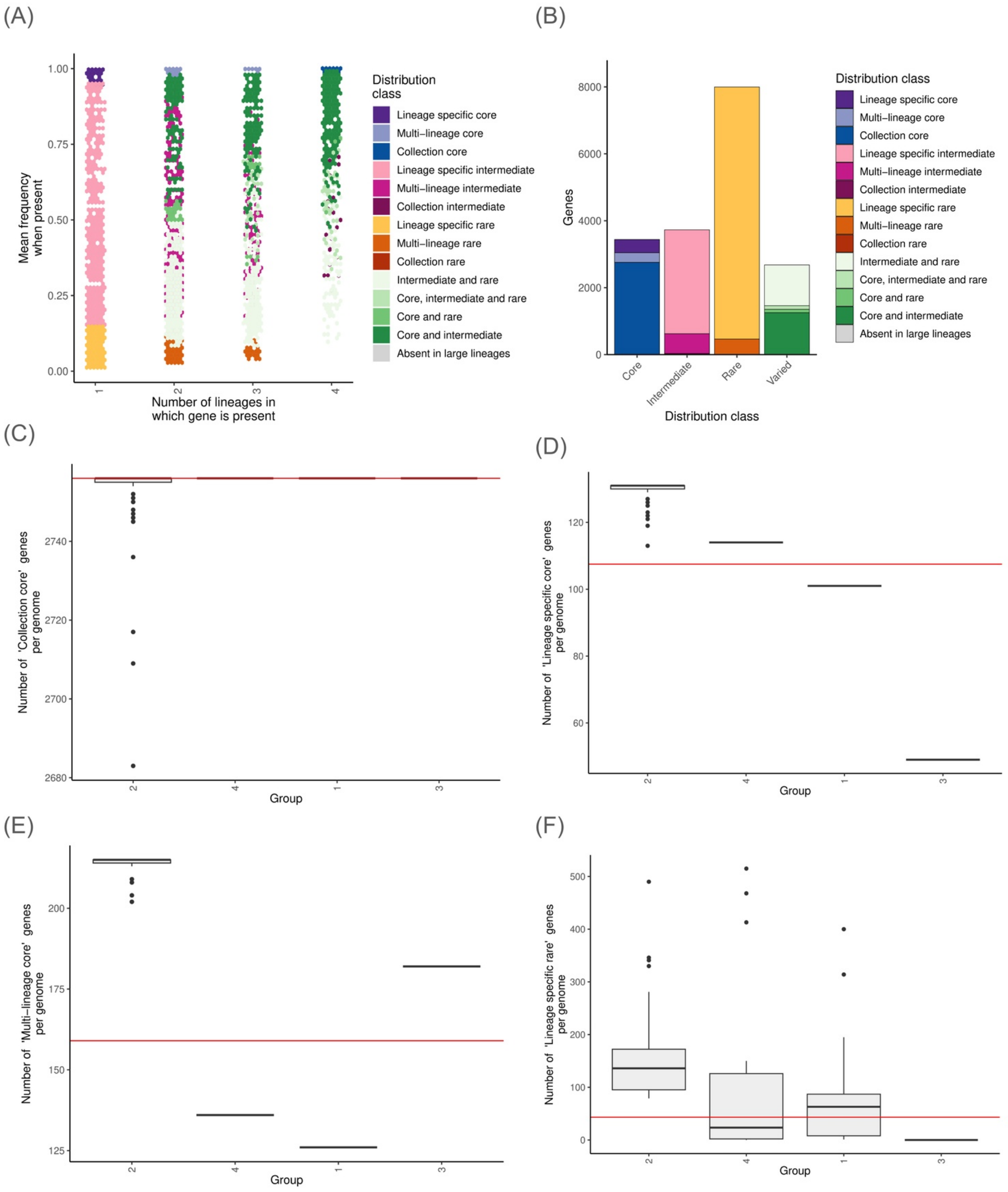
(A)
Mean frequency when present
Number of lineages in which gene is present
Distribution class
Lineage specific core
Multi−lineage core
Collection core
Lineage specific intermediate
Multi−lineage intermediate
Collection intermediate
Lineage specific rare
Multi−lineage rare
Collection rare
Intermediate and rare
Core, intermediate and rare
Core and rare
Core and intermediate
Absent in large lineages
(B)
Genes
Core
Intermediate
Rare
Varied
Distribution class
(C)
Number of 'Collection core' genes per genome
Group
(D)
Number of 'Lineage specific core' genes per genome
Group
(E)
Number of 'Multi−lineage core' genes per genome
Group
(F)
Number of 'Lineage specific rare' genes per genome
Group

**Supplementary Figure 3. Relative frequency of twilight pangenome partition distribution classes in different RSSC lineages.** **(A)** Relative frequency representation as Hexbin plot, **(B)** Number of genes in each twilight distribution class. Distribution of four differenct twilight distribution class for aggregated for each lineage: **(C)** Collection core, **(D)** lineage specific core, **(E)** Multi-lineage core, **(F)** Lineage specific rare

48

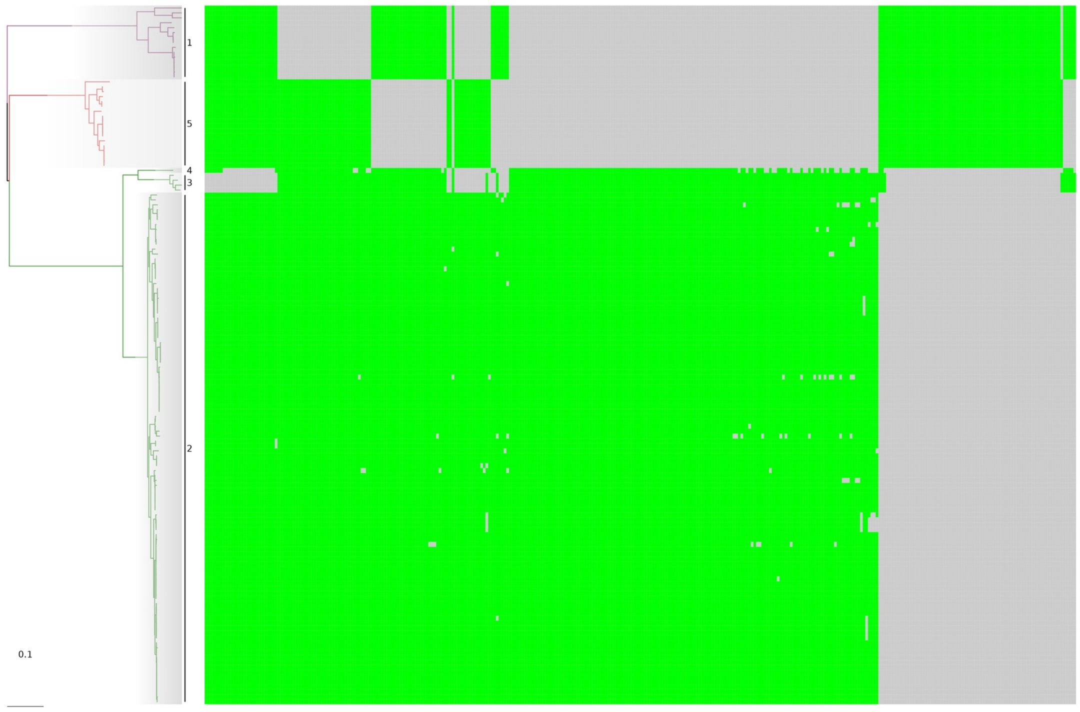

**Supplementary Figure 4. Multi-lineage core genes in RSSC phylogeny and population structure.** Columns are genes and rows indicate isolates placed in the tree from core genome phylogeny. Green color indicates presence while grey colors denotes absense of genes in a given isolates.

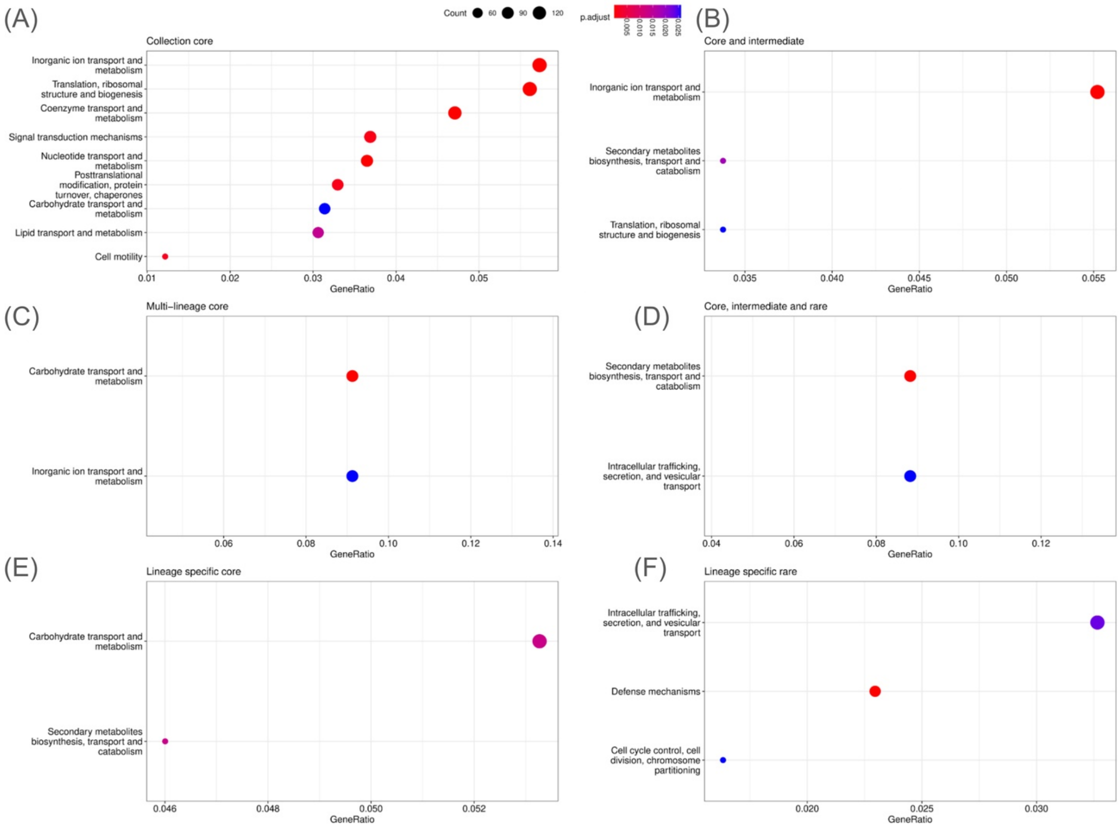

(A)
Collection core
Inorganic ion transport and metabolism
Translation, ribosomal structure and biogenesis
Coenzyme transport and metabolism
Signal transduction mechanisms
Nucleotide transport and metabolism
Posttranslational modification, protein turnover, chaperones
Carbohydrate transport and metabolism
Lipid transport and metabolism
Cell motility
0.01
0.02
0.03
0.04
0.05
GeneRatio
Count
60
90
120
p.adjust
0.005
0.010
0.015
0.020
0.025
(B)
Core and intermediate
Inorganic ion transport and metabolism
Secondary metabolites biosynthesis, transport and catabolism
Translation, ribosomal structure and biogenesis
0.035
0.040
0.045
0.050
0.055
GeneRatio
(C)
Multi−lineage core
Carbohydrate transport and metabolism
Inorganic ion transport and metabolism
0.06
0.08
0.10
0.12
0.14
GeneRatio
(D)
Core, intermediate and rare
Secondary metabolites biosynthesis, transport and catabolism
Intracellular trafficking, secretion, and vesicular transport
0.04
0.06
0.08
0.10
0.12
GeneRatio
(E)
Lineage specific core
Carbohydrate transport and metabolism
Secondary metabolites biosynthesis, transport and catabolism
0.046
0.048
0.050
0.052
GeneRatio
(F)
Lineage specific rare
Intracellular trafficking, secretion, and vesicular transport
Defense mechanisms
Cell cycle control, cell division, chromosome partitioning
0.020
0.025
0.030
GeneRatio

**Supplementary Figure 5. COG enrichment analysis across different twilight pangenome categories.** **(A)** Collection core, **(B)** Core and intermediate, **(C)** Multi-lineage core, **(D)** Core, intermediate, rare, **(E)** Lineage specific core, **(F)** Lineage specific rare.

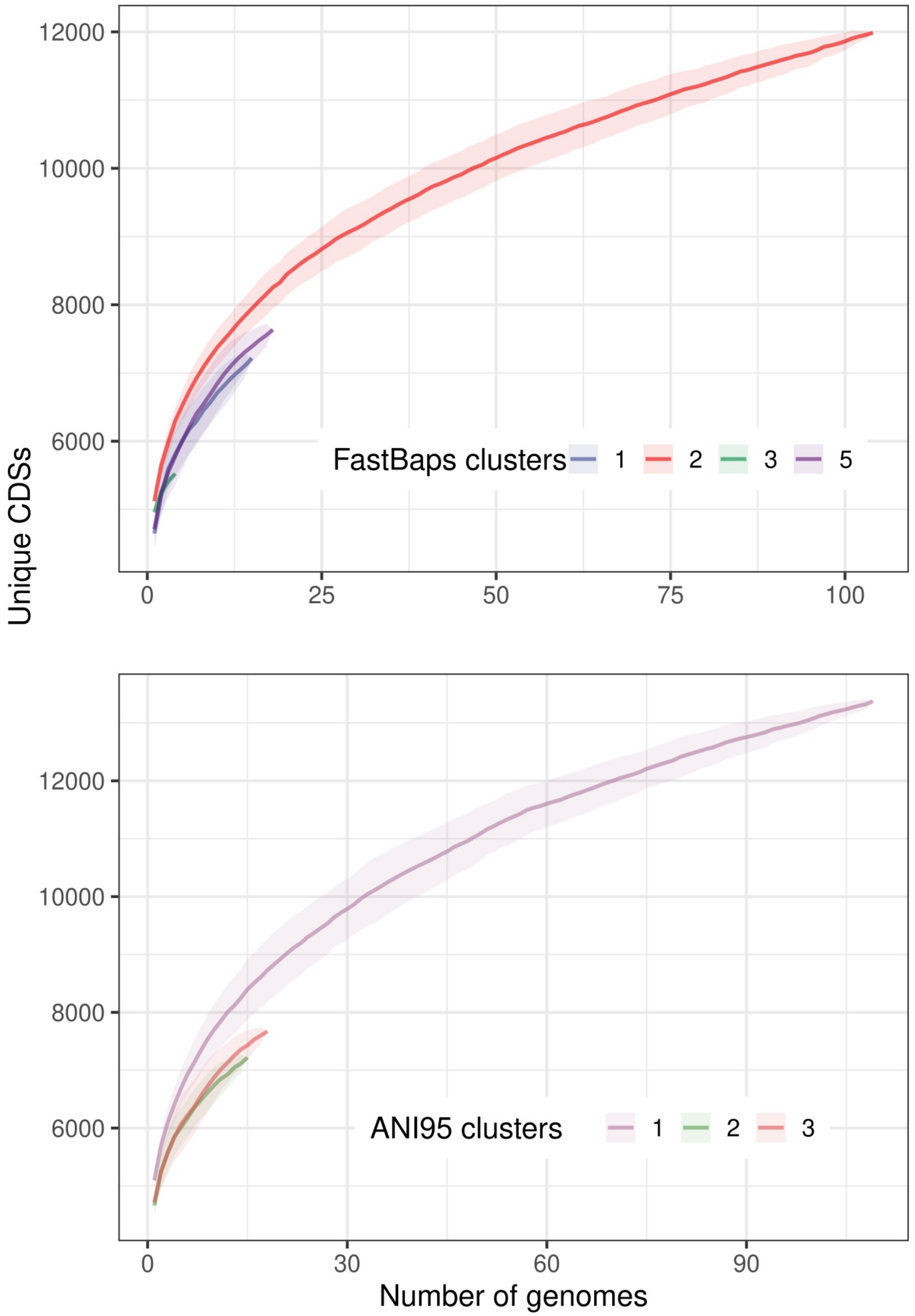

Unique CDSs
12000
10000
8000
6000
FastBaps clusters
1
2
3
5
0
25
50
75
100
12000
10000
8000
6000
ANI95 clusters
1
2
3
0
30
60
90
Number of genomes

**Supplementary Figure 6. Pangenome expansion dynamics and phylogenetic structure.** Gene accumulation curves demonstrate the open nature of the RSSC pangenome across 142 complete genomes. (Top) Rarefaction curves colored by FastBAPS population structure clusters (1-5), showing differential gene accumulation rates among phylogenetic lineages. Cluster 2 (red) exhibits the steepest accumulation curve, indicating greater genetic diversity and larger accessory genome content. (Bottom) Curves stratified by ANI-defined species boundaries (*R. solanacearum*, *R. pseudosolanacearum*, *R. syzygii*), revealing species-specific pangenome expansion patterns. The consistently upward trajectory of all curves and Heaps' law alpha parameter of 0.58 ($< 1$) confirms an open pangenome where novel genes continue to be discovered with each additional genome sequenced, indicating ongoing horizontal gene transfer and adaptive evolution within the RSSC complex.

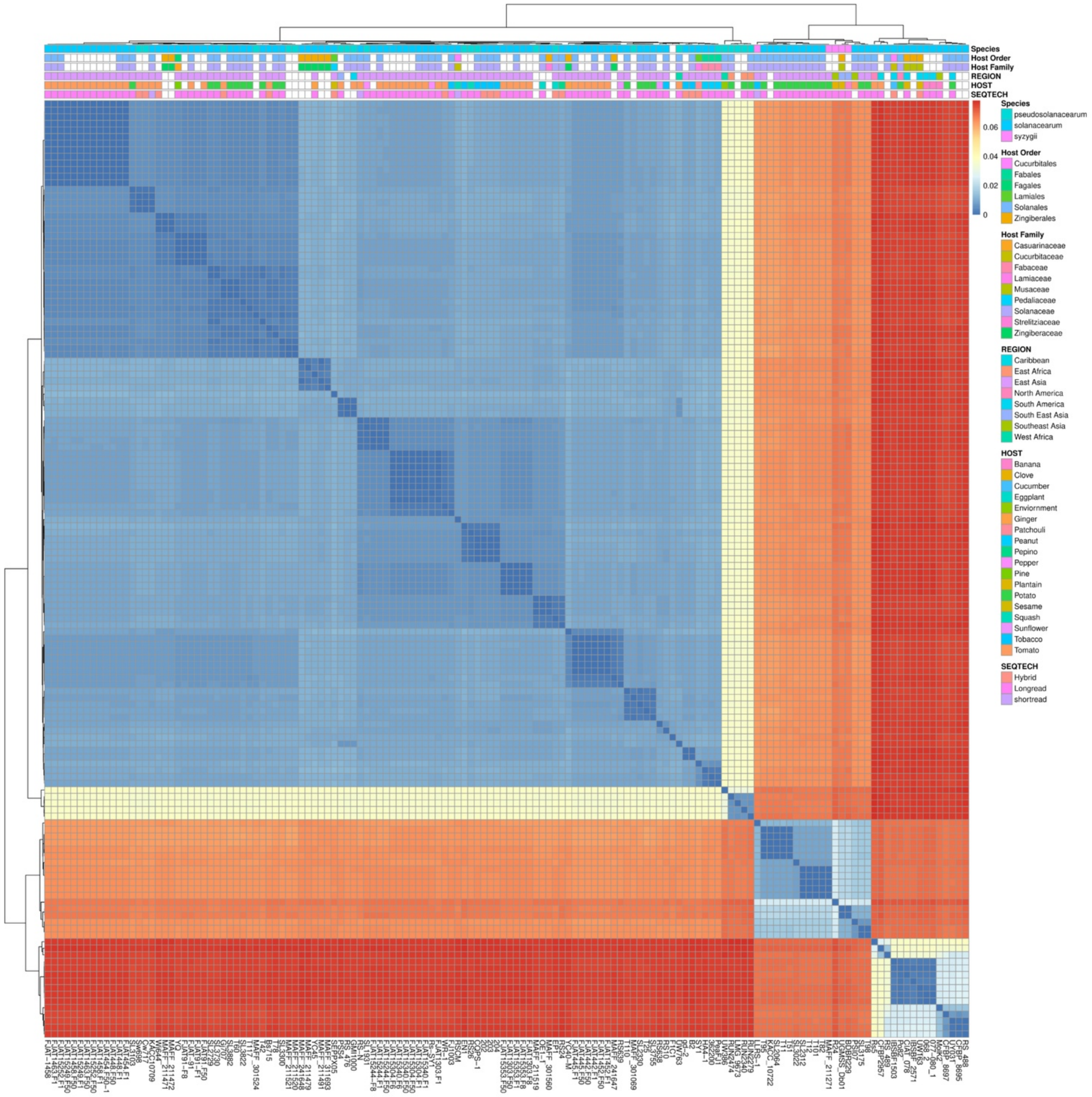
Species
Host Order
Host Family
REGION
HOST
SEQTECH
0.06
0.04
0.02
0
Species
pseudosolanacearum
solanacearum
syzygii
Host Order
Cucurbitales
Fabales
Fagales
Lamiales
Solanales
Zingiberales
Host Family
Casuarinaceae
Cucurbitaceae
Fabaceae
Lamiaceae
Musaceae
Pedaliaceae
Solanaceae
Strelitziaceae
Zingiberaceae
REGION
Caribbean
East Africa
East Asia
North America
South America
South East Asia
Southeast Asia
West Africa
HOST
Banana
Clove
Cucumber
Eggplant
Enviornment
Ginger
Patchouli
Peanut
Pepino
Pepper
Pine
Plantain
Potato
Sesame
Squash
Sunflower
Tobacco
Tomato
SEQTECH
Hybrid
Longread
shortread

**Supplementary Figure 7. Population genetic structure revealed by accessory genome architecture.** Hierarchical clustering heatmap based on pairwise Jaccard similarity coefficients calculated from binary accessory gene presence/absence profiles across 142 RSSC genomes. The color intensity represents similarity values (blue = high similarity, red = low similarity), with clear phylogenetic signal evident in the block-diagonal structure. The clustering pattern faithfully recapitulates the established phylotypic classification (I–IV) while revealing previously unrecognized substructure within Phylotype IV, specifically distinguishing Solanaceae-associated strains from Musaceae-associated strains. This fine-scale ecological differentiation pattern demonstrates that accessory genome content, rather than core genome phylogeny, is the primary driver of host-specific adaptation and niche specialization in the RSSC complex.

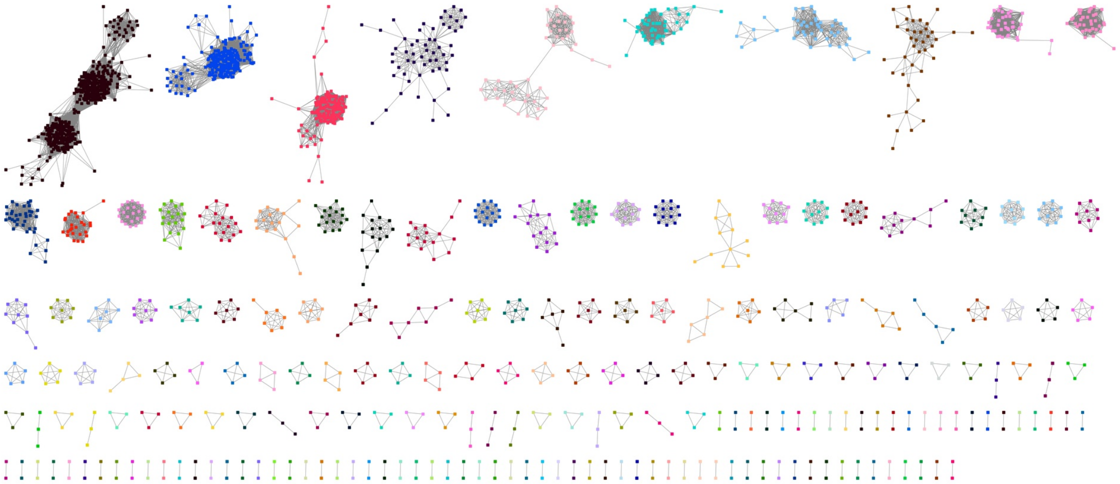

82

**Supplementary Figure 8. Co-incident accessory genesets network in RSSC that are significantly enriched than random chances.** Genes are represented as nodes and two genes(nodes) are connected by an edge if those genes are statistically associated, size of the node is weighted for lineage independence, nodes are colored by connected-component of the graph which indicate associated gene sets.

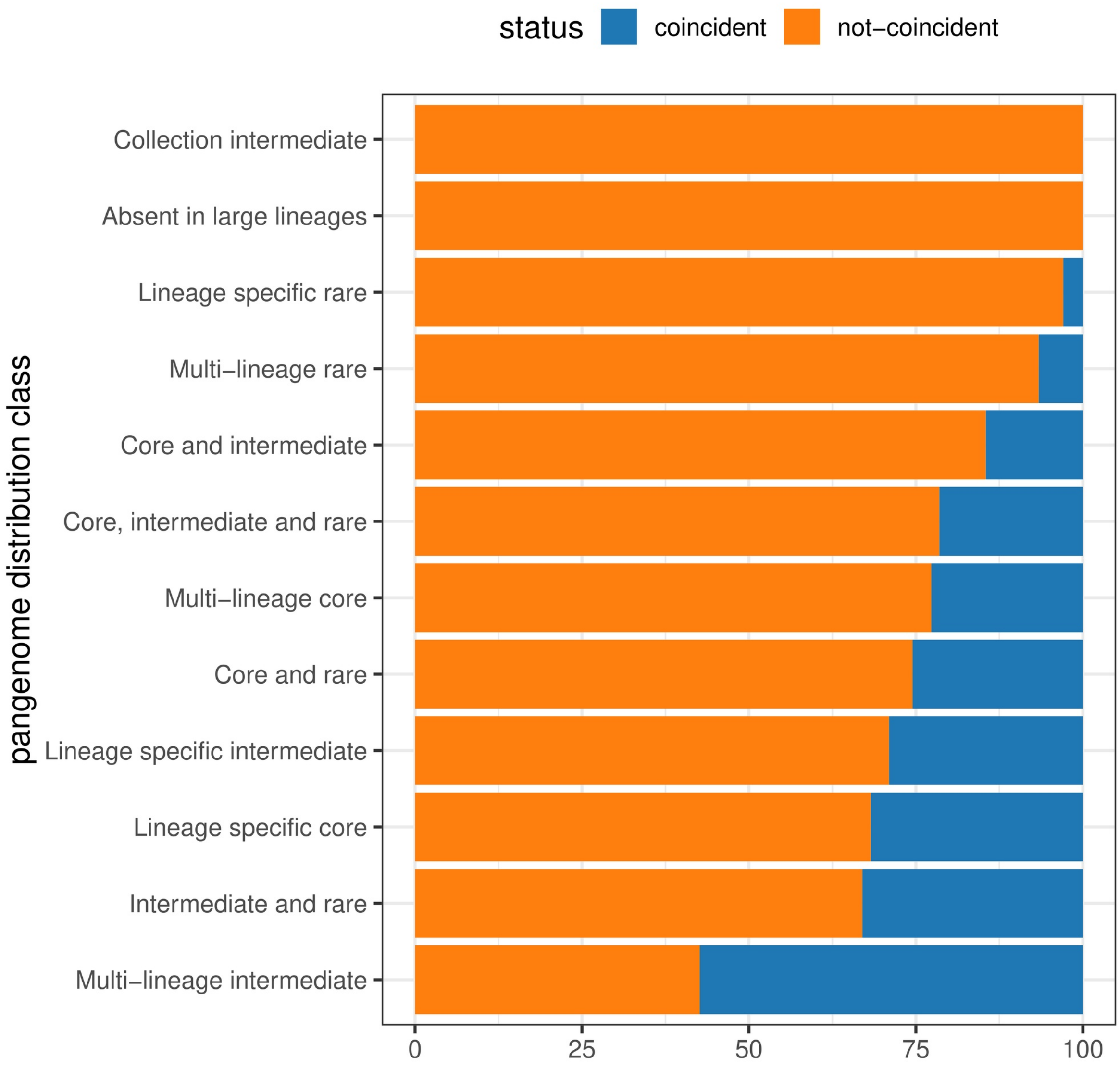
status
coincident
not−coincident
pangenome distribution class
Collection intermediate
Absent in large lineages
Lineage specific rare
Multi−lineage rare
Core and intermediate
Core, intermediate and rare
Multi−lineage core
Core and rare
Lineage specific intermediate
Lineage specific core
Intermediate and rare
Multi−lineage intermediate
0
25
50
75
100

**Supplementary Figure 9. Significantly associated co-incident accessory genes in different twilight distribution classes.**

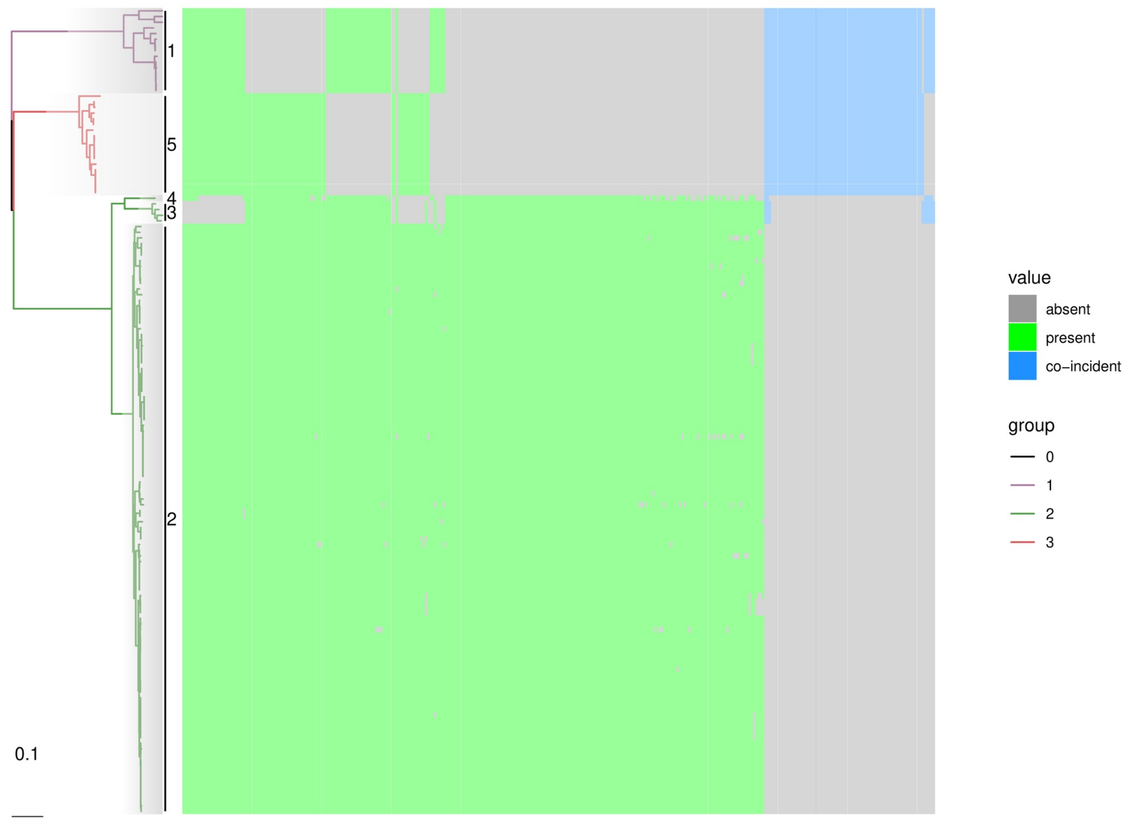
1
5
4
3
2
0.1
value
absent
present
co−incident
group
0
1
2
3

**Supplementary Figure 10. Co-incident genes in Multi-lineage core as shown in previous diagram, here co-incident genes are highlighted in blue.**

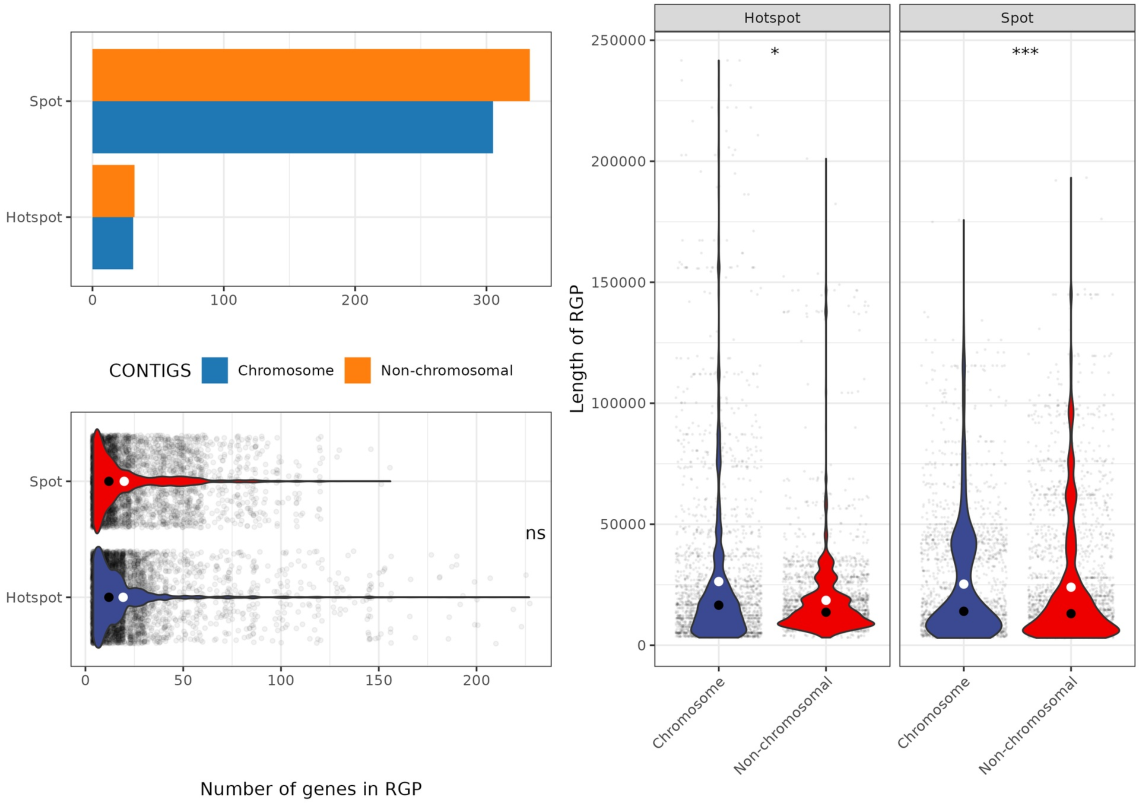
Spot
Hotspot
0
100
200
300
CONTIGS
Chromosome
Non-chromosomal
ns
50
150
Number of genes in RGP
Hotspot
Spot
*
***
250000
200000
150000
100000
50000
Length of RGP
Chromosome
Non-chromosomal

**Supplementary Figure 11. Structural and distributional characteristics of genomic plasticity regions.** Comprehensive analysis of Regions of Genome Plasticity (RGPs) organization and integration patterns. (Top left) Bar chart showing the distribution of genomic spots between chromosomal (n=327) and non-chromosomal replicons (n=324, including megaplasmids and smaller plasmids), indicating approximately equal spot numbers but different functional specializations. (Bottom left) Gene density comparison between standard spots (<100 genes) and gene-rich hotspots (>100 genes), with hotspots representing sites of extensive horizontal gene transfer and genetic innovation. (Right) Violin plots comparing RGP length distributions between replicon types, demonstrating that chromosomal RGPs tend to be longer and more variable in size compared to plasmid-borne RGPs. Wilcoxon rank-sum statistical tests confirm significant differences in length distributions between replicon types ($p < 0.05$, $^{**}p < 0.001$), suggesting distinct evolutionary pressures and integration mechanisms for chromosomal versus extrachromosomal genetic elements.

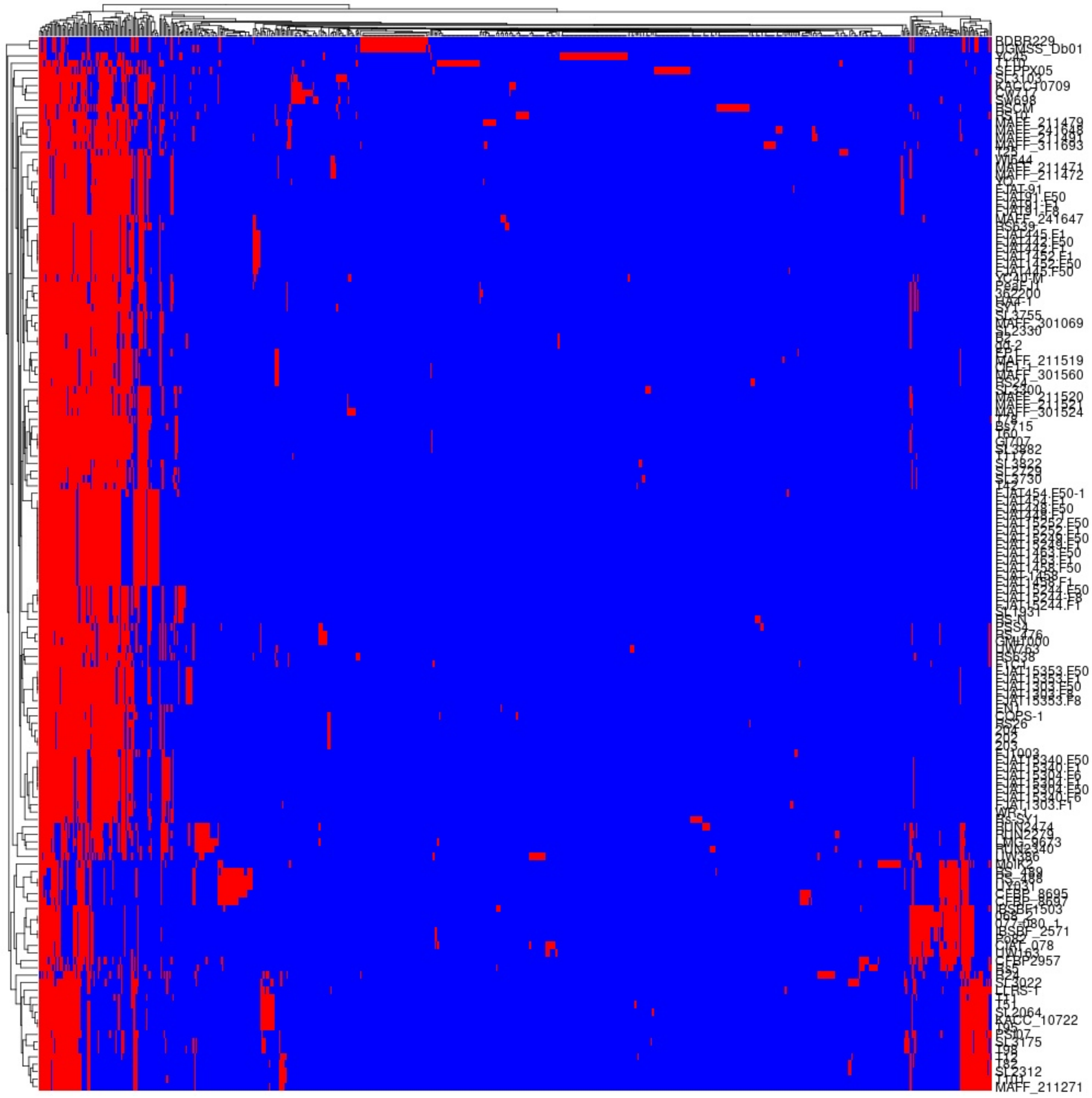

**Supplementary Figure 12. Genomic spot utilization patterns across the RSSC population.** Binary presence/absence matrix illustrating the contribution of individual isolates (rows, n=142) to specific genomic integration spots (columns, n=651). Red cells indicate isolate contribution to spot gene content, while blue cells represent absence of contribution. The matrix reveals distinct patterns: (1) ubiquitous spots present across most isolates, likely representing core accessory functions; (2) lineage-specific spots restricted to phylogenetically related strains; (3) rare spots found in only a few isolates, potentially representing recent acquisition events or specialized adaptations. The hierarchical clustering of both isolates and spots reveals the modular organization of the accessory genome, where certain spot combinations co-occur, suggesting functional linkage or shared evolutionary origins. This pattern underlies the flexibility of RSSC genomes in adapting to diverse ecological niches through differential accessory gene content.

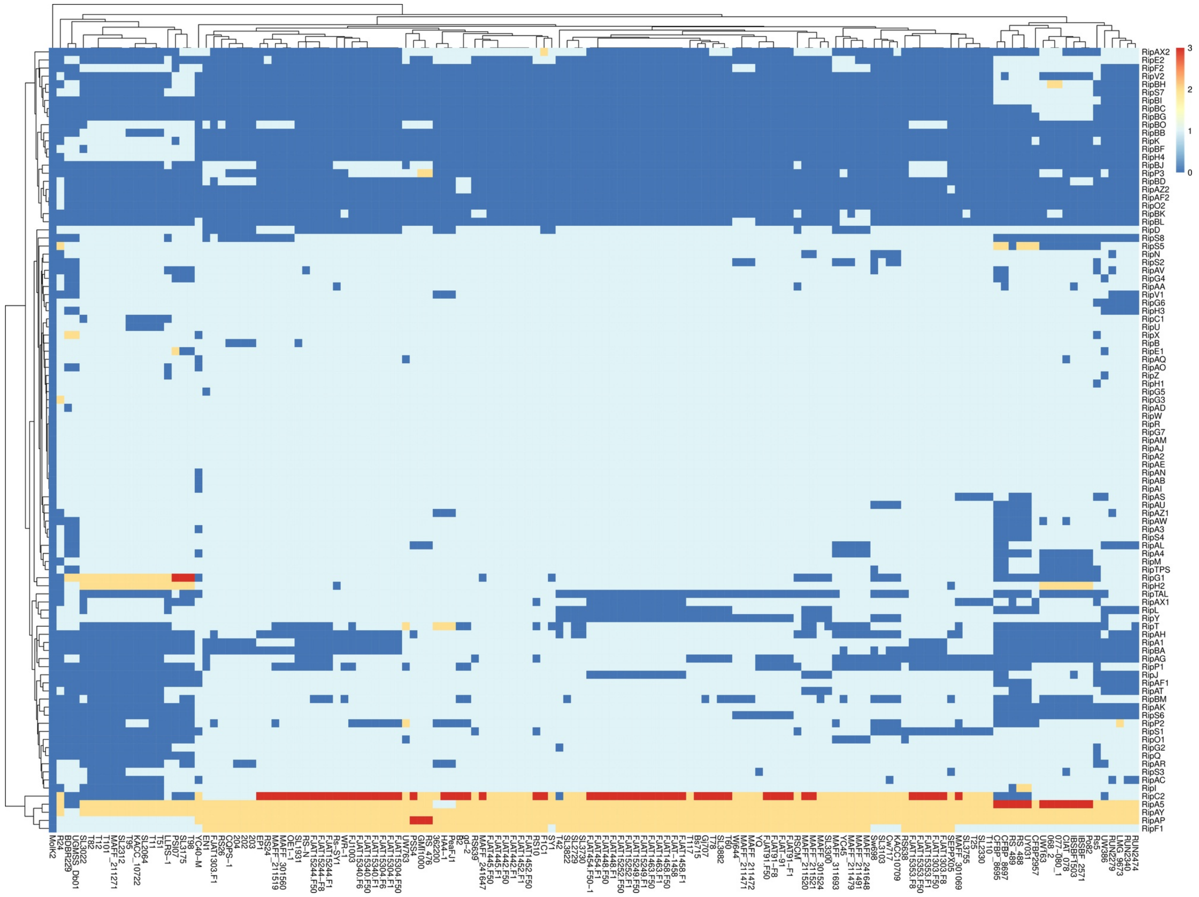
3
2
1
0
RipAX2
RipE2
RipF2
RipV2
RipBH
RipS7
RipBI
RipBC
RipBG
RipBO
RipBB
RipK
RipBF
RipH4
RipBJ
RipP3
RipBD
RipAZ2
RipAF2
RipO2
RipBK
RipBL
RipD
RipS8
RipS5
RipN
RipS2
RipAV
RipG4
RipAA
RipV1
RipG6
RipH3
RipC1
RipU
RipX
RipB
RipE1
RipAQ
RipAO
RipZ
RipH1
RipG5
RipG3
RipAD
RipW
RipR
RipG7
RipAM
RipAJ
RipA2
RipAE
RipAN
RipAB
RipAI
RipAS
RipAU
RipAZ1
RipAW
RipA3
RipS4
RipAL
RipA4
RipM
RipTPS
RipG1
RipH2
RipTAL
RipAX1
RipL
RipY
RipT
RipAH
RipA1
RipBA
RipAG
RipP1
RipJ
RipAF1
RipAT
RipBM
RipAK
RipS6
RipP2
RipS1
RipO1
RipG2
RipQ
RipAR
RipS3
RipAC
RipI
RipC2
RipA5
RipAY
RipAP
RipF1

**Supplementary Figure 13. Modular organization of Type III secretion system effector repertoires across genomic integration sites.** Comprehensive heatmap showing the presence/absence distribution of T3SS effector genes (Rip proteins) across genomic spots in the RSSC population. Rows represent individual effector genes (n=95 different Rip types), columns represent genomic spots, and colored cells indicate effector detection within specific integration loci. The analysis reveals that 50% of T3SS genes (5,495 out of 10,853 total secretory genes) are incorporated into spots, with distinct modular organization patterns: (1) spot-specific effector modules where subsets of effectors consistently co-occur (e.g., RipH3 concentrated in spot_164, RipAN in spot_16); (2) distributed effectors found across multiple spots, potentially reflecting different evolutionary origins; (3) lineage-specific effector combinations that correspond to host adaptation strategies. This modular architecture indicates that virulence repertoires are assembled through discrete genome plasticity events rather than gradual accumulation, with specific spots serving as "virulence cassettes" that can be horizontally transferred as functional units.

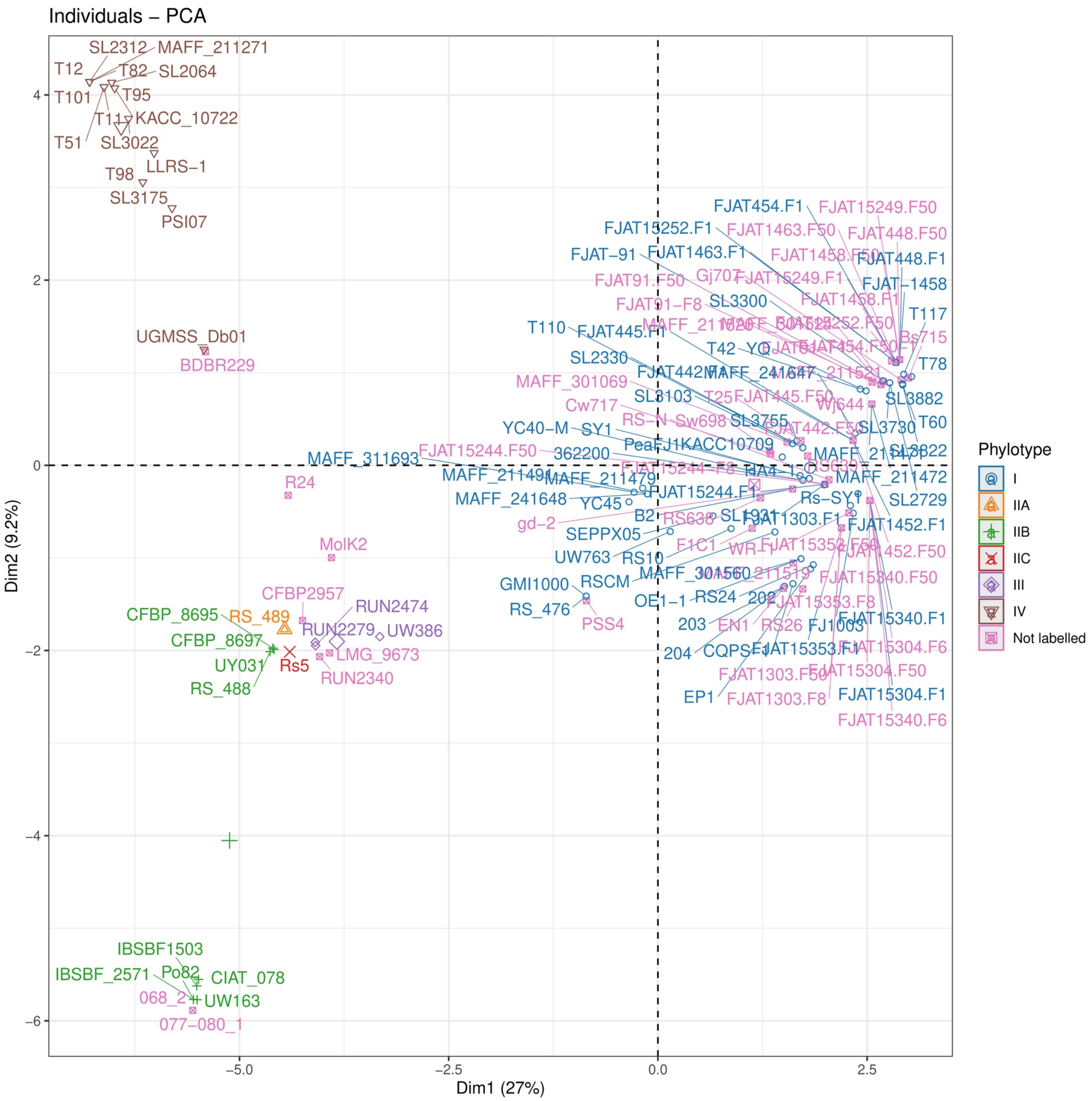
Individuals – PCA
Dim1 (27%)
Dim2 (9.2%)
Phylotype
I
IIA
IIB
IIC
III
IV
Not labelled
SL2312
MAFF_211271
T12
T82
SL2064
T101
T95
T11
KACC_10722
T51
SL3022
LLRS–1
T98
SL3175
PSI07
UGMSS_Db01
BDBR229
R24
MolK2
CFBP2957
RUN2474
CFBP_8695
RS_489
RUN2279
UW386
CFBP_8697
LMG_9673
UY031
Rs5
RUN2340
RS_488
IBSBF1503
IBSBF_2571
Po82
CIAT_078
068_2
UW163
077–080_1
MAFF_311693
FJAT15244.F50
MAFF_241648
gd–2
GMI1000
RS_476
PSS4
FJAT454.F1
FJAT15249.F50
FJAT448.F50
FJAT448.F1
FJAT–1458
T117
T78
SL3882
T60
SL2729
FJAT1452.F1
FJAT15340.F50
FJAT15340.F1
FJAT15304.F50
FJAT15304.F1
FJAT15340.F6
EP1
FJAT1303.F8
204
203
RS24
4
2
0
–2
–4
–6
–5.0
–2.5
0.0
2.5

**Supplementary Figure 14. Phylogenetic constraint on effector repertoire diversification revealed by multivariate analysis.** Principal component analysis (PCA) of RSSC isolates based on binary T3SS effector gene presence/absence profiles across 95 different Rip protein families. Each point represents an individual isolate projected onto the first two principal components, with colors denoting phylotype classification (I–IV, plus unlabeled isolates). The clear separation of phylotype clusters demonstrates that effector repertoires are strongly constrained by phylogenetic history, with PC1 and PC2 together explaining approximately 30% of the total variance. Notable patterns include: (1) tight clustering within phylotypes, indicating conserved core effector sets; (2) phylotype-specific regions of PC space, reflecting lineage-associated effector innovations; (3) intermediate positions for some isolates, suggesting horizontal transfer events or hybrid repertoires. This ordination confirms that despite the modular organization of effectors in genomic spots, major effector repertoire differences align with established phylogenetic boundaries, indicating that host adaptation occurs within the framework of vertical inheritance.

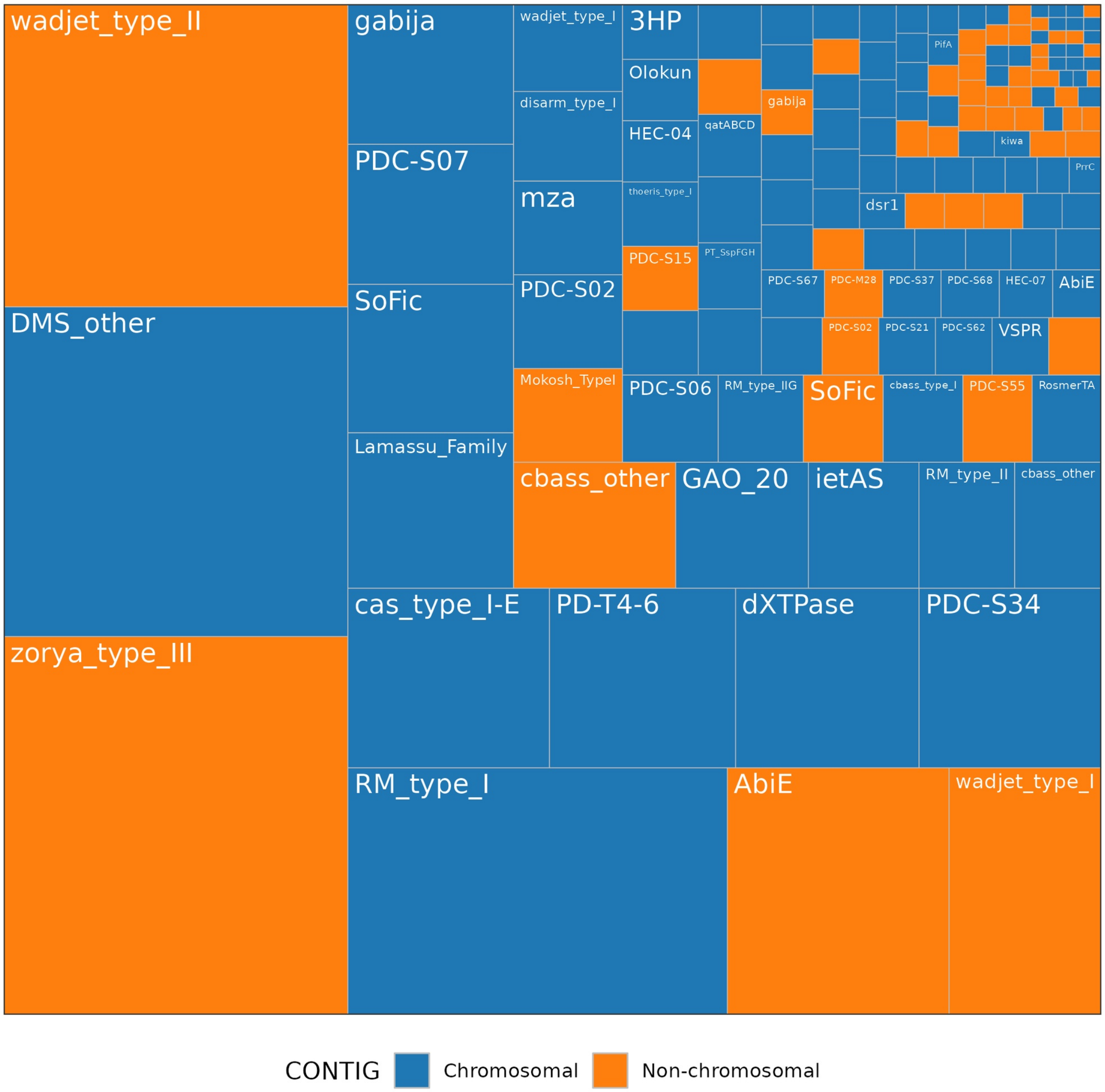
wadjet_type_II
DMS_other
zorya_type_III
gabija
PDC-S07
SoFic
Lamassu_Family
wadjet_type_I
disarm_type_I
mza
PDC-S02
Mokosh_TypeI
3HP
Olokun
HEC-04
thoeris_type_I
PDC-S15
qatABCD
PT_SspFGH
gabija
PifA
kiwa
PrrC
dsr1
PDC-S67
PDC-M28
PDC-S37
PDC-S68
HEC-07
AbiE
PDC-S02
PDC-S21
PDC-S62
VSPR
PDC-S06
RM_type_IIG
SoFic
cbass_type_I
PDC-S55
RosmerTA
cbass_other
GAO_20
ietAS
RM_type_II
cbass_other
cas_type_I-E
PD-T4-6
dXTPase
PDC-S34
RM_type_I
AbiE
wadjet_type_I
CONTIG
Chromosomal
Non-chromosomal

**Supplementary Figure 15. Replicon-specific integration preferences of bacterial defense systems.** Treemap visualization showing the relative abundance of defense systems types identified by PADLOC across 142 RSSC genomes. Rectangle sizes are proportional to system abundance, with colors indicating chromosomal (blue) or non-chromosomal/plasmid (orange) localization preferences. Major systems like zorya_type_III, RM_type_I, and DMS_other show predominantly chromosomal integration, reflecting their essential roles in cellular defense and stable inheritance. Conversely, systems such as wadjet_type_II, gabija, and AbiE demonstrate marked preference for plasmid integration, suggesting more recent acquisition, horizontal transfer potential, and possible role in rapid adaptation to phage pressure. This differential distribution pattern indicates that defense system evolution in RSSC involves both stable chromosomal systems and mobile plasmid-borne defenses that can spread rapidly through populations.

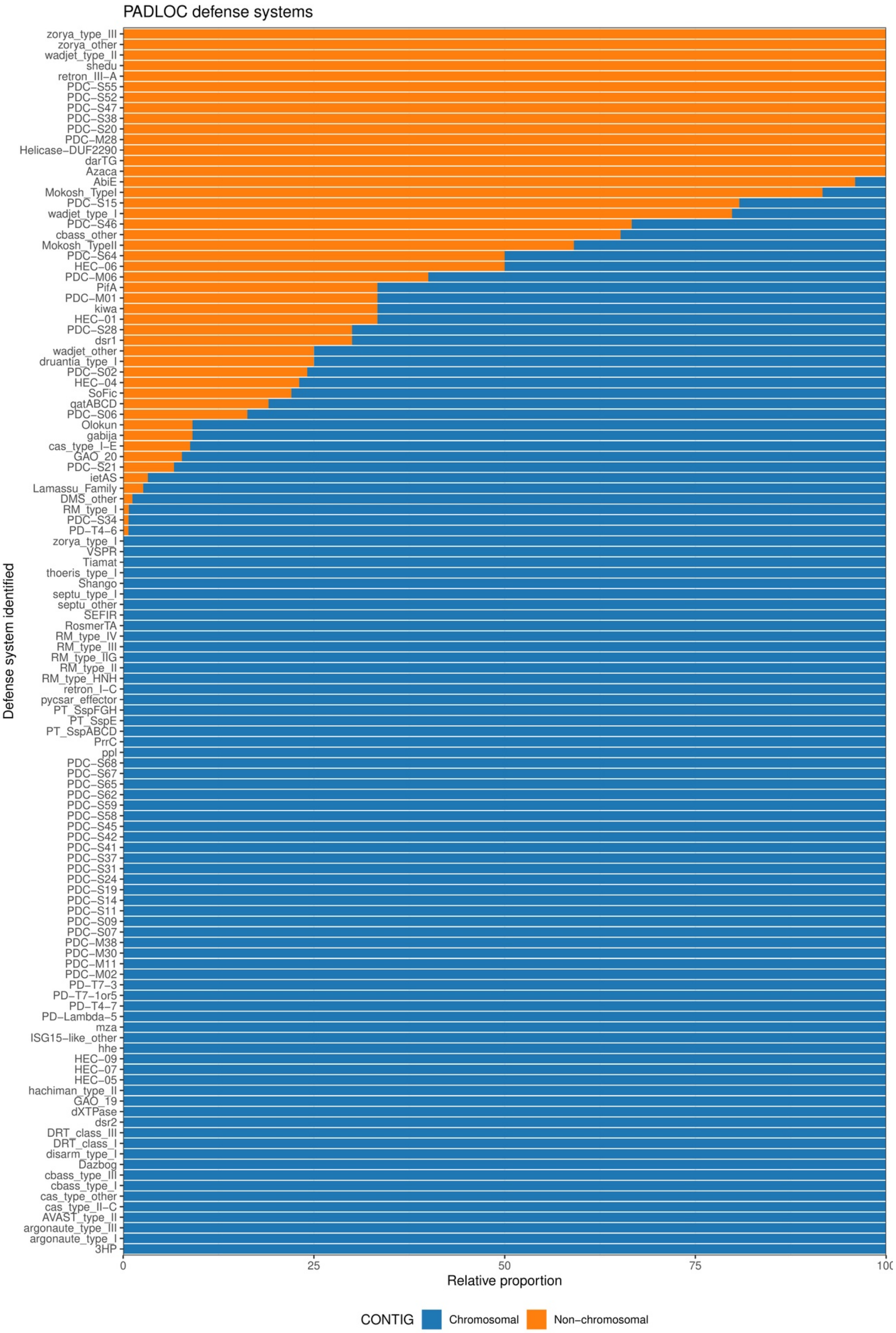
PADLOC defense systems
zorya_type_III
zorya_other
wadjet_type_II
shedu
retron_III−A
PDC−S55
PDC−S52
PDC−S47
PDC−S38
PDC−S20
PDC−M28
Helicase−DUF2290
darTG
Azaca
AbiE
Mokosh_TypeI
PDC−S15
wadjet_type_I
PDC−S46
cbass_other
Mokosh_TypeII
PDC−S64
HEC−06
PDC−M06
PifA
PDC−M01
kiwa
HEC−01
PDC−S28
dsr1
wadjet_other
druantia_type_I
PDC−S02
HEC−04
SoFic
qatABCD
PDC−S06
Olokun
gabija
cas_type_I−E
GAO_20
PDC−S21
ietAS
Lamassu_Family
DMS_other
RM_type_I
PDC−S34
PD−T4−6
zorya_type_I
VSPR
Tiamat
thoeris_type_I
Shango
septu_type_I
septu_other
SEFIR
RosmerTA
RM_type_IV
RM_type_III
RM_type_IIG
RM_type_II
RM_type_HNH
retron_I−C
pycsar_effector
PT_SspFGH
PT_SspE
PT_SspABCD
PrrC
ppl
PDC−S68
PDC−S67
PDC−S65
PDC−S62
PDC−S59
PDC−S58
PDC−S45
PDC−S42
PDC−S41
PDC−S37
PDC−S31
PDC−S24
PDC−S19
PDC−S14
PDC−S11
PDC−S09
PDC−S07
PDC−M38
PDC−M30
PDC−M11
PDC−M02
PD−T7−3
PD−T7−1or5
PD−T4−7
PD−Lambda−5
mza
ISG15−like_other
hhe
HEC−09
HEC−07
HEC−05
hachiman_type_II
GAO_19
dXTPase
dsr2
DRT_class_III
DRT_class_I
disarm_type_I
Dazbog
cbass_type_III
cbass_type_I
cas_type_other
cas_type_II−C
AVAST_type_II
argonaute_type_III
argonaute_type_I
3HP
Defense system identified
0
25
50
75
100
Relative proportion
CONTIG
Chromosomal
Non−chromosomal

**Supplementary Figure 16. Relative proportion of each defense system in Chromosomal vs Non-chromosomal contigs (Megaplasmids and other contigs) characterized by PADLOC**

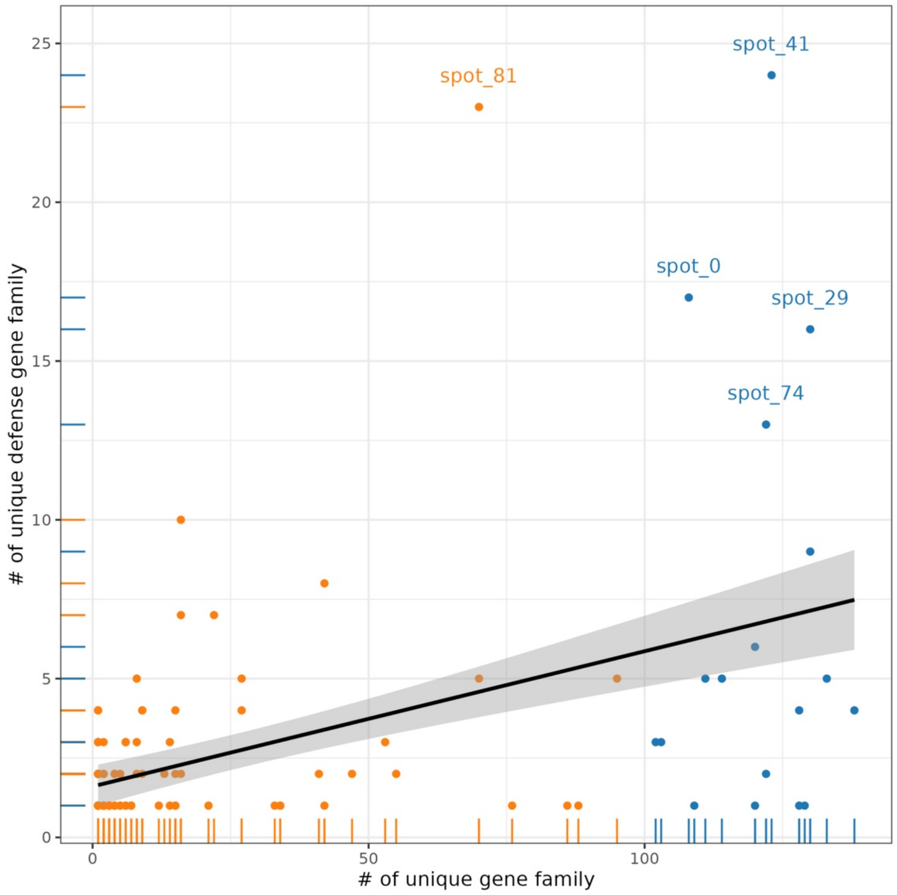
spot_41
spot_81
spot_0
spot_29
spot_74
### of unique defense gene family
### of unique gene family
0
5
10
15
20
25
0
50
100

168

**Supplementary Figure 17. Relation of number of gene-families in defensive systems and total number of unique gene families in spot.** Orange colored points indicate spots with less than 100 unique gene family while blue spots are spots with more than 100 gene families, also referred to hotspots.

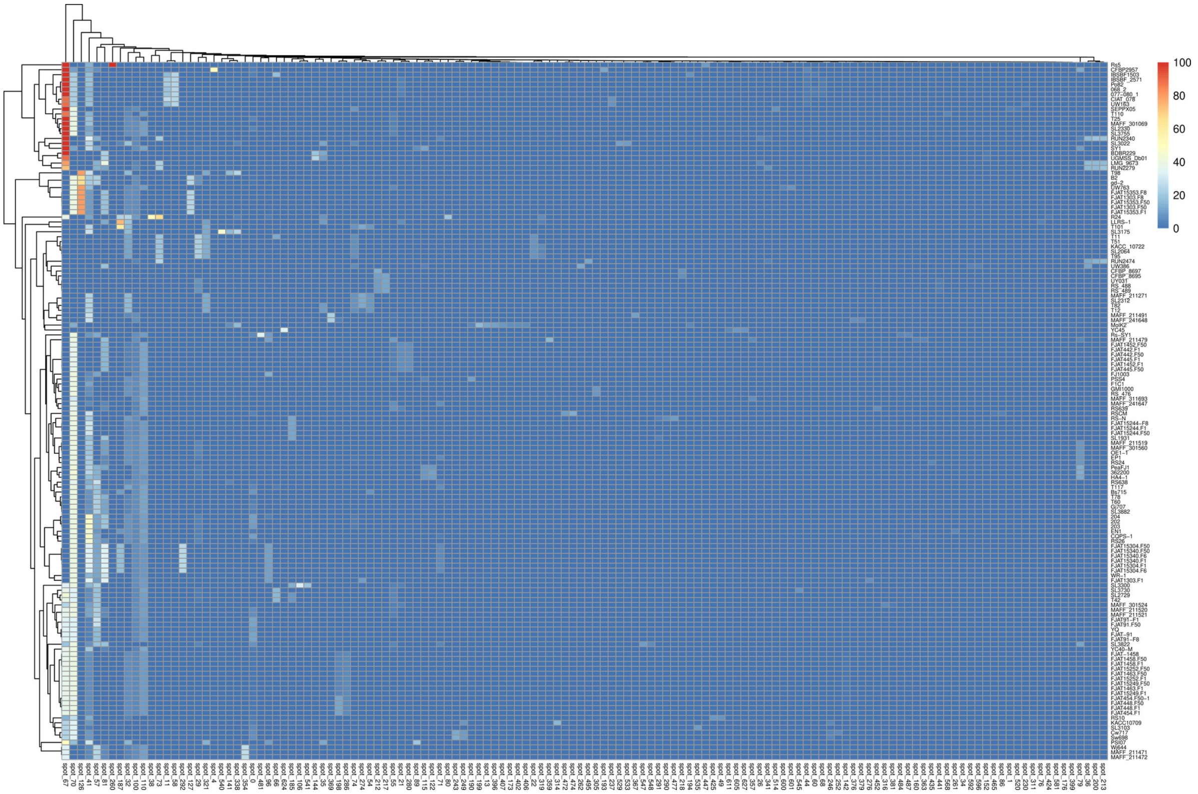
100
80
60
40
20
0

**Supplementary Figure 18. Spatial organization of defense genes within genomic plasticity regions.** Intensity heatmap displaying defense gene density within specific genomic spots across the 142-genome RSSC collection. Each cell represents the number of PADLOC-identified defense genes present in a given spot (columns) for each isolate (rows), with color intensity proportional to gene count (blue = absent, orange/red = high density). The non-random distribution reveals that 73% of all defense genes (3,840 out of 4,870 total) are concentrated within discrete plasticity regions rather than being uniformly distributed across genomes. Notable patterns include: (1) defense "hotspots" where multiple systems co-localize, (2) isolate-specific defense profiles reflecting different phage pressures, and (3) coordinated presence/absence of entire defense gene clusters, suggesting these regions function as integrated defense modules that are gained or lost as units during evolution.

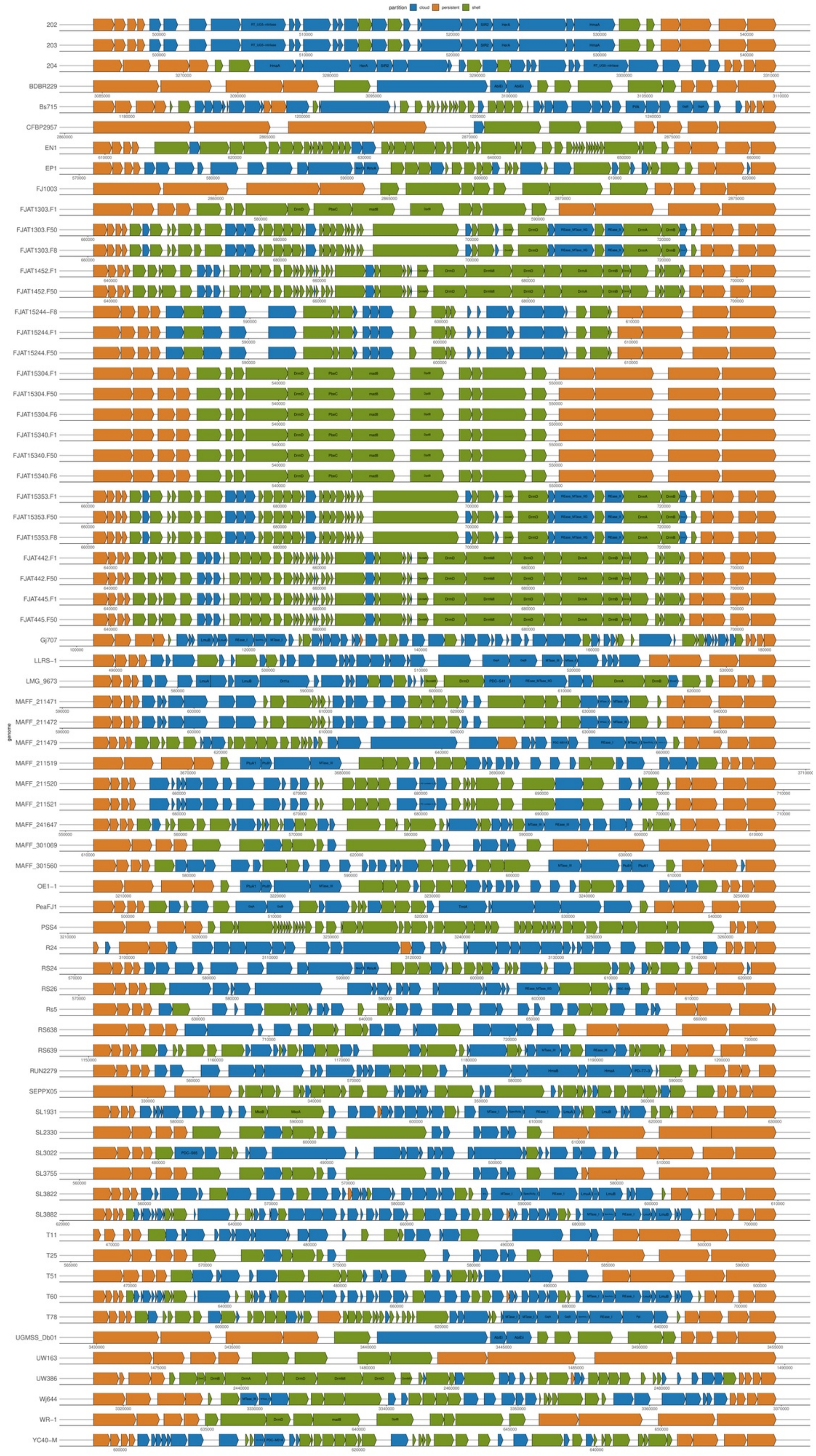
partition
cloud
persistent
shell
genome
202
203
204
BDBR229
Bs715
CFBP2957
EN1
EP1
FJ1003
FJAT1303.F1
FJAT1303.F50
FJAT1303.F8
FJAT1452.F1
FJAT1452.F50
FJAT15244-F8
FJAT15244.F1
FJAT15244.F50
FJAT15304.F1
FJAT15304.F50
FJAT15304.F6
FJAT15340.F1
FJAT15340.F50
FJAT15340.F6
FJAT15353.F1
FJAT15353.F50
FJAT15353.F8
FJAT442.F1
FJAT442.F50
FJAT445.F1
FJAT445.F50
Gj707
LLRS-1
LMG_9673
MAFF_211471
MAFF_211472
MAFF_211479
MAFF_211519
MAFF_211520
MAFF_211521
MAFF_241647
MAFF_301069
MAFF_301560
OE1-1
PeaFJ1
PSS4
R24
RS24
RS26
Rs5
RS638
RS639
RUN2279
SEPPX05
SL1931
SL2330
SL3022
SL3755
SL3822
SL3882
T11
T25
T51
T60
T78
UGMSS_Db01
UW163
UW386
Wj644
WR-1
YC40-M

**Supplementary Figure 19. Organization of accessory genes in one representative** *spot*#81 **identified from panRGP framework.** Persistent (core) genes are in the flanks (in orange block) while the accessory genes (cloud and shell) are present between the flanking persistent genes.
